# The unique genome architecture of the devastating West African cacao black pod pathogen *Phytophthora megakarya*

**DOI:** 10.1101/826636

**Authors:** Abraham Morales-Cruz, Shahin S. Ali, Andrea Minio, Rosa Figueroa-Balderas, Jadran F. García, Takao Kasuga, Alina S. Puig, Jean-Philippe Marelli, Bryan A. Bailey, Dario Cantu

## Abstract

*Phytophthora megakarya* (*Pmeg*) and *P. palmivora* (*Ppal*) are oomycete pathogens that cause black pod rot of cacao (*Theobroma cacao*), the most economically important disease on cacao globally. While *Ppal* is a cosmopolitan pathogen, *Pmeg*, which is more aggressive on cacao than *Ppal*, has been reported only in West and Central Africa where it has been spreading and devastating cacao farms since the 1950s. In this study, we reconstructed the complete diploid genomes of multiple isolates of both species using single-molecule sequencing. Thirty-one additional genotypes were sequenced to analyze inter- and intra-species genomic diversity. These resources make it possible to better understand the molecular basis of virulence differences in closely related and consequential pathogens and study their evolutionary history. The *Pmeg* genome is exceptionally large (222 Mbp) and nearly twice the size *Ppal* (135 Mbp) and most known *Phytophthora* species (∼100 Mbp on average). We show that the genomes of both species recently expanded by independent whole-genome duplications (WGD). WGD and the dramatic transposable element associated expansion of a few gene families led to the exceptionally large genome and transcriptome of *Pmeg* and the diversification of virulence-related genes including secreted RxLR effectors. Finally, this study provides evidence of adaptive evolution among well-known effectors and discusses the implications of effector expansion and diversification.

## INTRODUCTION

Cacao (*Theobroma cacao* L.) beans are the backbone of the global chocolate industry, which is valued at over 100 billion US Dollars annually (1). Cacao is also the major cash crop for millions of small holder farmers in the tropics with a total harvest of 5.2 million tons (FAOSTATS, 2017). However, global cacao production is threatened by multiple diseases that negatively impact yield and quality (2). Black pod rot (BPR) is responsible for more than half the total reported crop loss, destroying the equivalent of 0.87 million metric tons or 2 billion US dollars’ worth of dried cacao beans annually (**Fig. 1A**) (2). Black pod rot is caused by multiple *Phytophthora* species, among which *P. palmivora* (*Ppal*) is the most widespread and *P. megakarya* (*Pmeg*), currently confined to West and Central Africa, is the most destructive, causing up to 90% crop loss if not controlled (**Fig. 1B**)(3). In recent years, *Pmeg* has largely displaced *Ppal* as a major cause of BPR in some African nations (4).

**Fig. 1.**
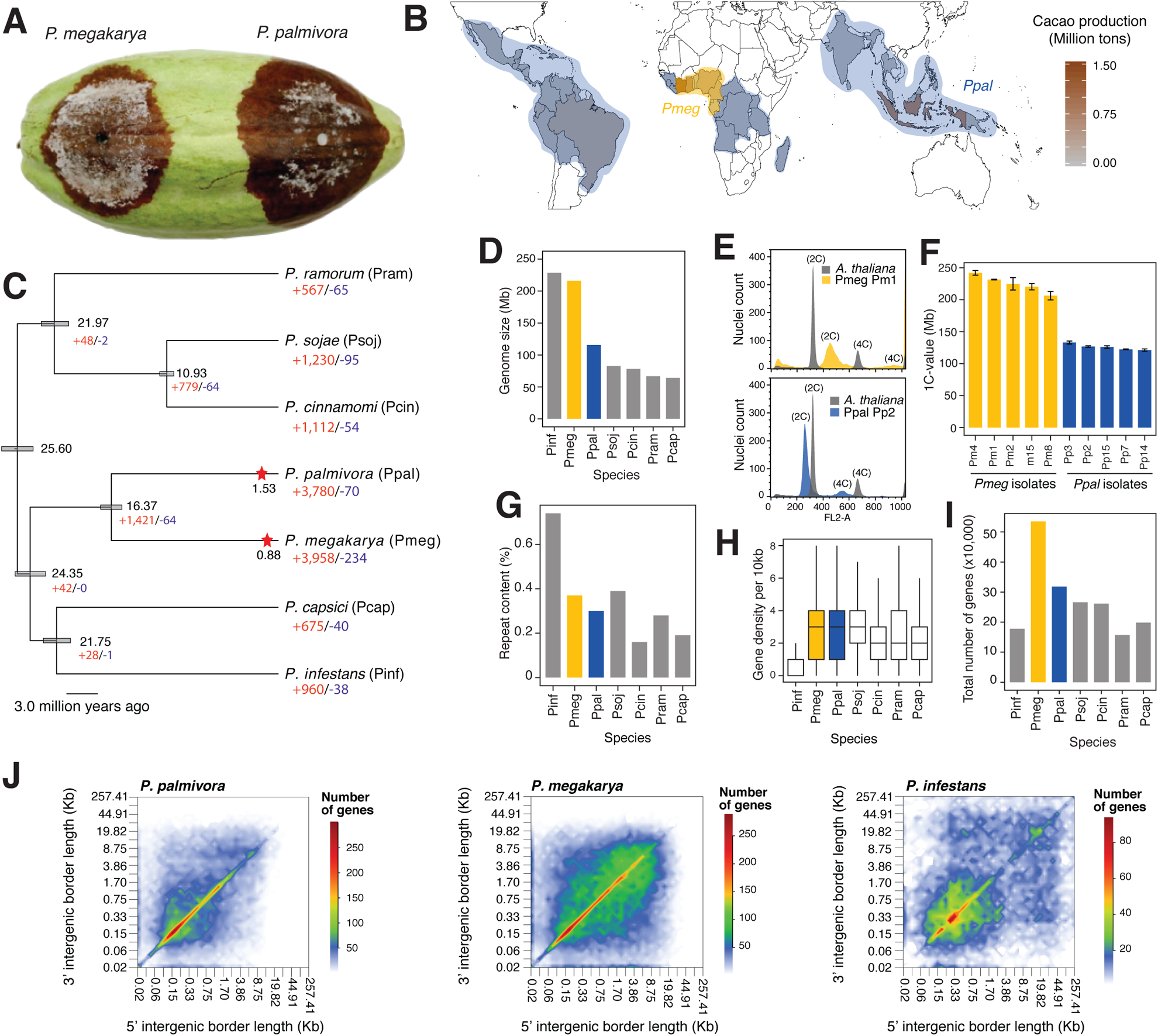
(A) Black pod rot symptoms by *Pmeg* and *Ppal.* (B) Approximate geographical distribution of *Pmeg* (yellow area) and *Ppal* (blue area). (C) Clock-calibrated phylogenetic tree with estimated times of divergence and WGD duplication events in *Ppal* and *Pmeg* in million years ago, with numbers of families expanded and contracted across multiple *Phytophtora* spp. (D) Genome size of multiple *Phytophthora* spp. (E) Example run of the flow cytometry using *Arabidopsis* as control. (F) Average estimated 1C-values from flow cytometry. (G) Repeat content of multiple *Phytophthora* spp. (H) Boxplot showing the distribution of gene density per 10 Kb. (I) Total number of genes per species. (J) Intergenic space heatmap of *Ppal, Pmeg* and *Pinf. Pinf*: *P. infestans, Pmeg*: *P. megakarya, Ppal*: *P. palmivora, Psoj*: *P. sojae, Pcin*: *P. cinnamomi, Pram*: *P. ramorum* and *Pcap*: *P. capsici.*

The genus *Phytophthora* includes more than 120 known species of filamentous oomycetes (5) and an estimated 300-500 species not yet discovered (6). Most of the known species are pathogenic to plants and some have devastating effects on crops and natural forests. There is expanding interest in *Phytophthora* species due to their economic and environmental impact. This has driven the release of genome assemblies for at least 26 *Phytophthora* species in the last 10 years. These genome assemblies have helped identify gene families critical to infection processes. These gene families include effectors like crinklers (CRNs), necrosis inducing proteins (NPPs), and proteins characterized by an N-terminal Arg-Xaa-Leu-Arg motif (RxLRs), all associated to pathogenicity and host specificity (7–9). Studies of *Phytophthora* genomes have shown that they contain significant amounts of repetitive elements. Transposable elements (TEs), specifically, play a significant role in the evolution of *Phytophthora* (4, 10–12). For example, in *P. ramorum* transposons have been associated with the generation of genotypic diversity by introducing chromosomal breakpoints (13).

Although *Pmeg* and *Ppal* are closely related and belong to clade 4 in the *Phytophthora* phylogeny (**Fig. 1C**) (14), they possess different numbers and sizes of chromosomes (9-12 smaller chromosomes in *Ppal*, 5-6 large chromosomes in *Pmeg*) and have different geographic distributions, host ranges, and aggressiveness (15, 16). Southeast Asia (17) and Central Africa (18) are the centers of origin of *Ppal* and *Pmeg*, respectively. Thus, they provide an opportunity to study the post-speciation divergent evolutionary trajectories of two pathogens that infect the same host with different levels of virulence. The interactions between each pathogen and cacao are likely to have been relatively recent. Initial draft genome assemblies for both species based on short-read sequencing suggested a whole-genome duplication (WGD) in *Ppal* and a retroelement-based expansion in the genome of *Pmeg*, resulting in particularly high numbers of RxLRs (4). However, complete genome assemblies for the two pathogens will be crucial to understanding their evolution and diversity and dissecting the mechanisms and adaptations that drive pathogenicity of BPR in cacao.

In this study, we assembled the diploid genomes of multiple isolates of *Pmeg* and *Ppal* using long reads. We found that the genome assembly size of *Pmeg* was nearly two-fold larger than previously reported (4), and validated the unexpected genome size by flow cytometry across multiple isolates. Further, we found strong evidence of WGDs in both species, as well as dramatic gene family expansions in *Pmeg* that likely explain the difference in the genomes of two species. We then examined highly expanded gene families in *Pmeg* and *Ppal* compared to other *Phytophthora* species, finding associations with transposable elements and large number of effectors. Finally, we re-sequenced 28 additional genotypes from cacao-producing countries and report evidence of adaptive evolution in well-known effectors that have recently increased in number by WGD and gene family expansion.

## RESULTS

### Phased assembly of long reads reveals the unique genome architecture of *P. megakarya*

Single molecule real-time long-read sequences were used to produce phased genome assemblies for three *P. megakarya* (*Pmeg*) and three *P. palmivora* (*Ppal*) isolates collected from cacao plantations in Western Africa (Pm1, Pm4, Pm15, Pp2), Central America (Pp3), and Southeast Asia (Pp15) (**Table 1**, SI Appendix, Tables S1-S3). All genome assemblies had a median coverage higher than 150x, high sequence accuracy (>99.999%), and gene space completeness comparable with other *Phytophthora* species assemblies (SI Appendix, Table S4).

**Table 1.** Genome assembly statistics for *P. megakarya* (*Pmeg*) and *P. palmivora* (*Ppal*). Average values ± SD are shown.

|  | *Pmeg* | *Ppal* |
| --- | --- | --- |
| Size (Mbp) | 222.04±25.19 | 135.32±17.21 |
| Number of contigs | 478.67±155.32 | 169.67±61.08 |
| N50 length (Mbp) | 1.02±0.56 | 1.53±0.56 |
| GC percentage | 0.49±0.01 | 0.49±0 |

The average sizes of the highly contiguous assemblies were 222.0 ± 25.2 Mbp and 135.3 ± 17.2 Mbp for *Pmeg* and *Ppal*, respectively (**Fig. 1D and Table 1**). Genome sizes of both species were confirmed using multiple assembly methods and flow cytometry (**Fig. 1E and F**; SI Appendix, Table S5 and S6). The differences between the haploid genome assembly sizes and the inferred 1C-values estimated by flow cytometry were less than 15% (*Pmeg*: 14.7 ± 5.6 Mb, *Ppal*: 13.5 ± 3.7 Mb). The similar values of the haploid genome assembly with the 1C-values validated the assembly sizes and suggests that both species are diploid (4). Flow cytometry confirmed the genome size in additional isolates of both species (**Fig. 1F**; SI Appendix, Table S5).

We then compared the *Ppal* and *Pmeg* genomes with previously published genomes of *Phytophthora* species (**Table 2**). Although *Pmeg* and *P. infestans* (*Pinf*) have similarly sized genomes, the structure of *Pmeg* was strikingly different from *Pinf* and any other *Phytophthora* genome. *Pinf*’s large genome size (229 Mb) was caused by the proliferation of TEs, which account for up to 70% of its genome (**Fig. 1D;** (8), which led to frequent gene-sparse regions with a large amount of intergenic space (**Fig. 1G**). Despite a larger genome, the amount of repetitive content in *Pmeg* (36.8 ± 1.2%) is similar to *Ppal* (33.5 ± 1.6%) and other *Phytophthora* spp. (25.5 ± 6.2%; **Fig. 1G**, SI Appendix, Table S7).

**Table 2.**
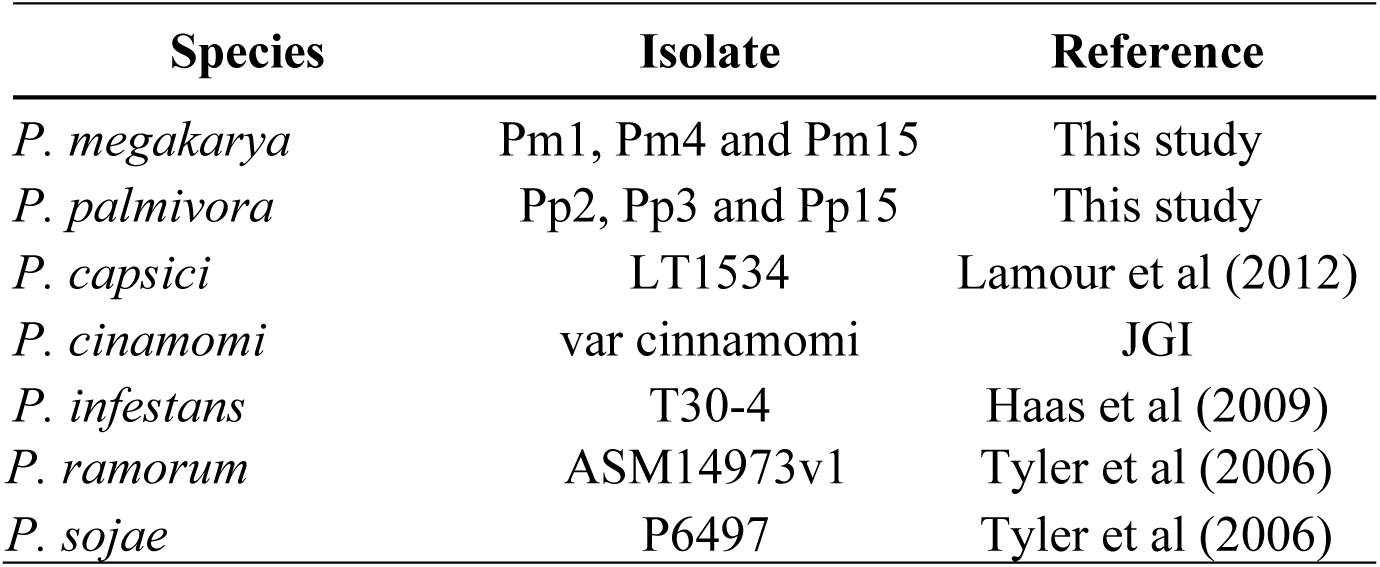
Genome assemblies analyzed in this study

Interestingly, the genomes of both cacao black pod pathogens have more protein-coding genes than other *Phytophthora* species. The primary assemblies of *Pmeg* and Ppal were predicted to have 57,577 ± 7,904 and 36,778 ± 4,481 protein-coding genes, respectively (**Fig. 1I**, Dataset S1). A high rate of duplication for eukaryotic universal single-copy orthologs (BUSCO) genes was observed; 36.7 ± 8.1% and 54.0 ± 11.3% of BUSCO genes were duplicated in *Pmeg* and *Ppal*, respectively. On average, only 11% of BUSCO genes were duplicated in other *Phytophthora* genomes analyzed using the same methods (SI Appendix, Table S4). The large percentage of duplicated conserved genes suggests that large-scale duplication processes occurred in the *Pmeg* and *Ppal* genomes. Like the small genomes of other *Phytophthora* spp. and in contrast to the gene-dispersed genome of *Pinf, Pmeg* has a high gene density despite its larger size (**Fig. 1J**). Overall, *Pmeg* has a highly dense genome with high frequency of genes in even the more dispersed sections of its genome and *Ppal* has a relatively compact genome, with most genes within 1 kbp of each other. Unlike *Pinf*, neither *Pmeg* nor *Ppal* contain highly expanded regions with intergenic spaces greater than 20 Kbp.

### Recent independent whole-genome duplications in *P. megakarya* and *P. palmivora*

The large number of genes in *Pmeg* and *Ppal* and the high rate of BUSCO gene duplication suggest a large-scale duplication process. Analysis of gene duplication across the entire gene space revealed 48,850 ± 9,475 (84.4 ± 4.5%) and 31,263 ± 6,015 (84.5 ± 6.6%) duplicated protein-coding genes in *Pmeg* and *Ppal*, respectively, compared to 10,744 ± 3,461 (49.87 ± 5.39%) duplicated genes in other *Phytophthora* genomes (**Fig. 2A**). We further classified duplications into the following patterns: conserved block duplicates (BD, minimum 5 genes), dispersed duplicates (DD), tandem duplicates (TD), genes in tandem and in block duplicates (BTD), and single-copy genes (SC). In both species, over 12,000 duplicated genes were organized in colinear BD genes (**Fig. 2A**), with each block including a median of 8 genes, but the two species greatly differed in the number of DD genes (**Fig. 2A**). Over 20,000 DD genes, about 40% of its entire transcriptome, were found in *Pmeg*. In contrast, ∼8,000 DD genes, only 25% its transcriptome, were found in *Ppal*. In both *Ppal* and *Pmeg*, duplicated blocks were often on the same scaffolds and one hundred kbp long, on average. Importantly, the presence of colinear duplicated blocks on the same scaffolds found both in the primary assembly and the haplotigs, confirmed the diploidy of the two species and ruled out colinear blocks belonging to homologous chromosomes (SI Appendix, Fig. S1). BD genes were virtually absent in the other *Phytophthora* genomes studied, with a maximum of only 476 BD genes in *Pinf.*

**Fig. 2.**
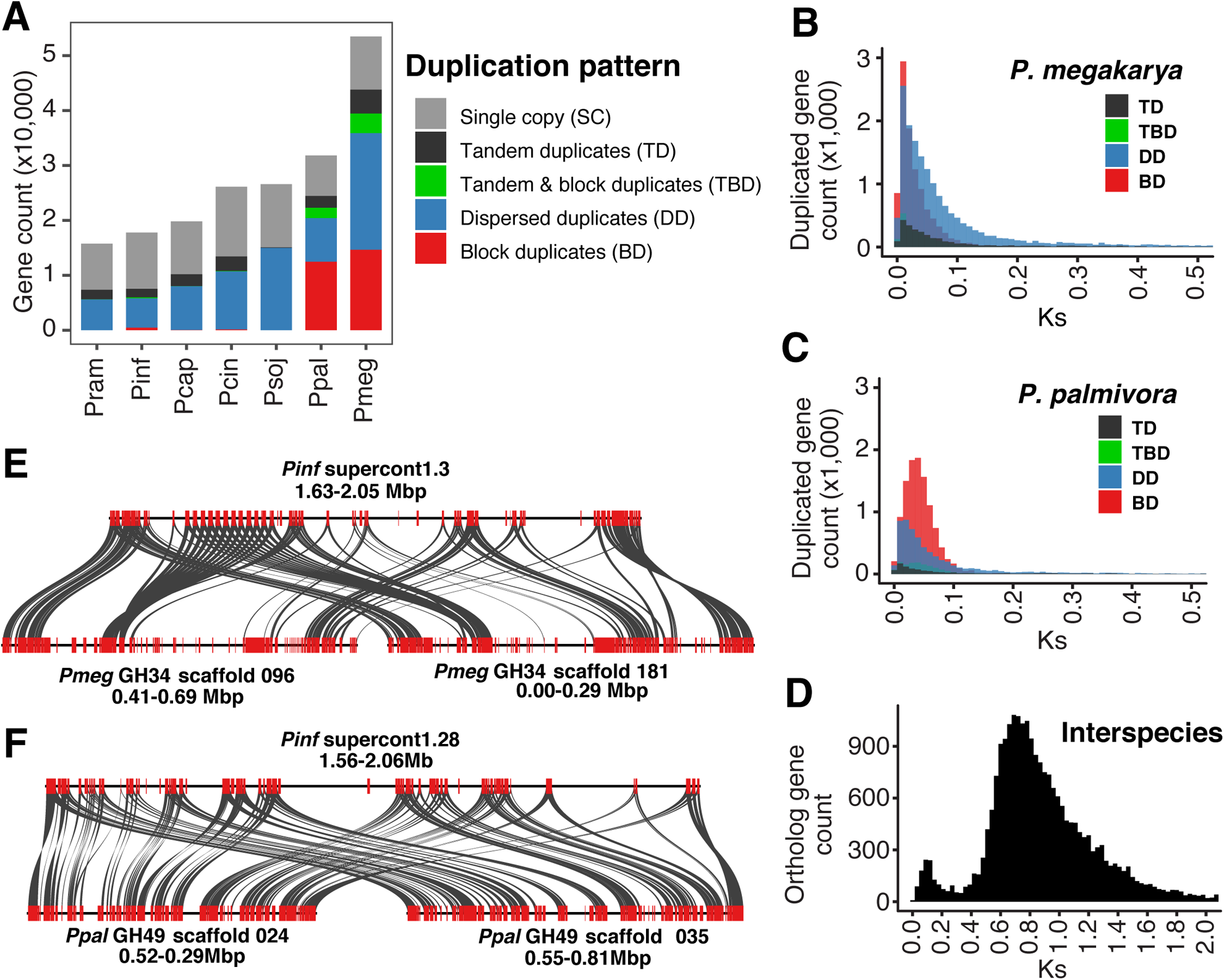
(A) Gene count per species divided by detected duplication patterns or the single copy category. Plots showing the synonymous mutations (Ks) of paralogs in *Pmeg* (B) and *Ppal* (C). Synteny of duplicated blocks in *Pmeg* (E) and in *Ppal* (F) with only one corresponding block in *Pinf*. (D) Plot showing orthologs Ks distribution between *Pmeg* and *Ppal*.

The large number of colinear BD genes suggests that a recent whole-genome duplication (WGD) happened in the two cacao pathogens. The inference of WGD in *Pmeg* and *Ppal* was supported by the synonymous substitutions rate (Ks) distribution across paralogous genes (**Fig. 2B&C**). Well-defined peaks in the Ks distributions were evident in both species indicating a sudden increase in new genes for *Pmeg* (0.021 Ks) and for *Ppal* (0.040 Ks) (**Fig. 2B&C**). The large-scale duplication events appear to have happened independently and after speciation (0.820 Ks; **Fig. 2D**). To estimate when the WGDs occurred, we searched for single-copy genes in other *Phytophthora* species that were present as two copies in *Pmeg* and *Ppal*. Of these, only the genes in conserved colinear blocks were considered further, since they were likely generated during a WGD event. By this standard, a WGD occurred 1.53 million years ago (MYA) in *Ppal* and 0.88 MYA in *Pmeg* (**Fig. 1C**, SI Appendix, Fig. S2). Since the divergence of *Pmeg* and *Ppal* was estimated at ∼16.4 MYA (**Fig. 1C**), we conclude that the WGD events happened independently and after the two species diverged.

Nearly 60% of all the genes were duplicated as part of the large-scale duplication process (<0.5 Ks) in both species, representing approximately 75% of the gene families in both species (SI Appendix, Fig. S3 and Dataset S1). In addition, several complete duplicated blocks in *Pmeg* and *Ppal* were colinear with a single block in *Pinf* (**Fig. 2E&F**; SI Appendix, Fig. S4). These results are strong evidence that independent WGD events occurred in both *Ppal* and *Pmeg.*

### Extensive gene expansion and duplicate dispersion in *P. megakarya*

Our analyses show that both *Pmeg* and *Ppal* have experienced recent WGD events, but these events do not explain their 40% difference in annotated protein coding genes. To better understand this difference, we focused on the number and organization of gene families across multiple *Phytophthora* species (**Table 2**). Using published transcriptomes, we found nearly 3,000 more gene families expanded in *Pmeg* and *Ppal* than the median number of families expanded in other *Phytophthora* species (e.g., *Pinf*: 960; **Fig. 1C**). Though the number of expanded gene families was similar for the two cacao pathogens, the number of predicted genes gained was much larger in *Pmeg* (27,166) than in *Ppal* (9,546).

Considering the large number of gene families expanded in these two species, hereinafter we focused on the gene families with at least a two-fold increase when compared to the median of the other *Phytophthora* species and containing at least 10 genes. The majority of these families were in *Pmeg* with 168 families (including 9,210 genes), while *Ppal* only had 96 such families (including 3,404 genes). The difference in the number of genes in the > 2-fold expanded families between *Pmeg* and *Ppal* indicates that there was a unique expansion of relatively few gene families in *Pmeg*.

Next, we studied the contribution of the duplication patterns to the > 2-fold expanded families in *Pmeg* and *Ppal*. The BD, TD, TBD and SC categories were relatively similar in both species, only differing by a maximum of 898 genes. However, *Pmeg* had 3,341 more DD genes than Ppal, suggesting that DD genes contributed disproportionately to the expansion of genes families in *Pmeg* and to its total number of genes. The mechanism by which duplication may have occurred was examined next.

### Expanded gene families are generally associated with transposable elements

TEs have the ability to mediate gene duplications of the host genes and the formation of the new genes (19–21). The proliferation of Long Terminal Repeat (LTR) TEs was reported to be the main cause of the genome expansion in *Pinf*, contributing to regions in the genome where gene order was not conserved, and the expansion of intergenic space (8). LTRs account for the large majority of the repetitive elements in the genome in *Pmeg* (76.3%) and *Ppal* (75.0%) (Dataset S2). Consequently, we studied the relationship between duplication patterns and LTRs by calculating the distance between genes and these elements (**Fig. 3A**). We found that DD genes were significantly closer (Kolmogorov-Smirnov test, *P-value <* 2.2e-16) to LTRs than to any other duplication category of genes in both species.

**Fig. 3.**
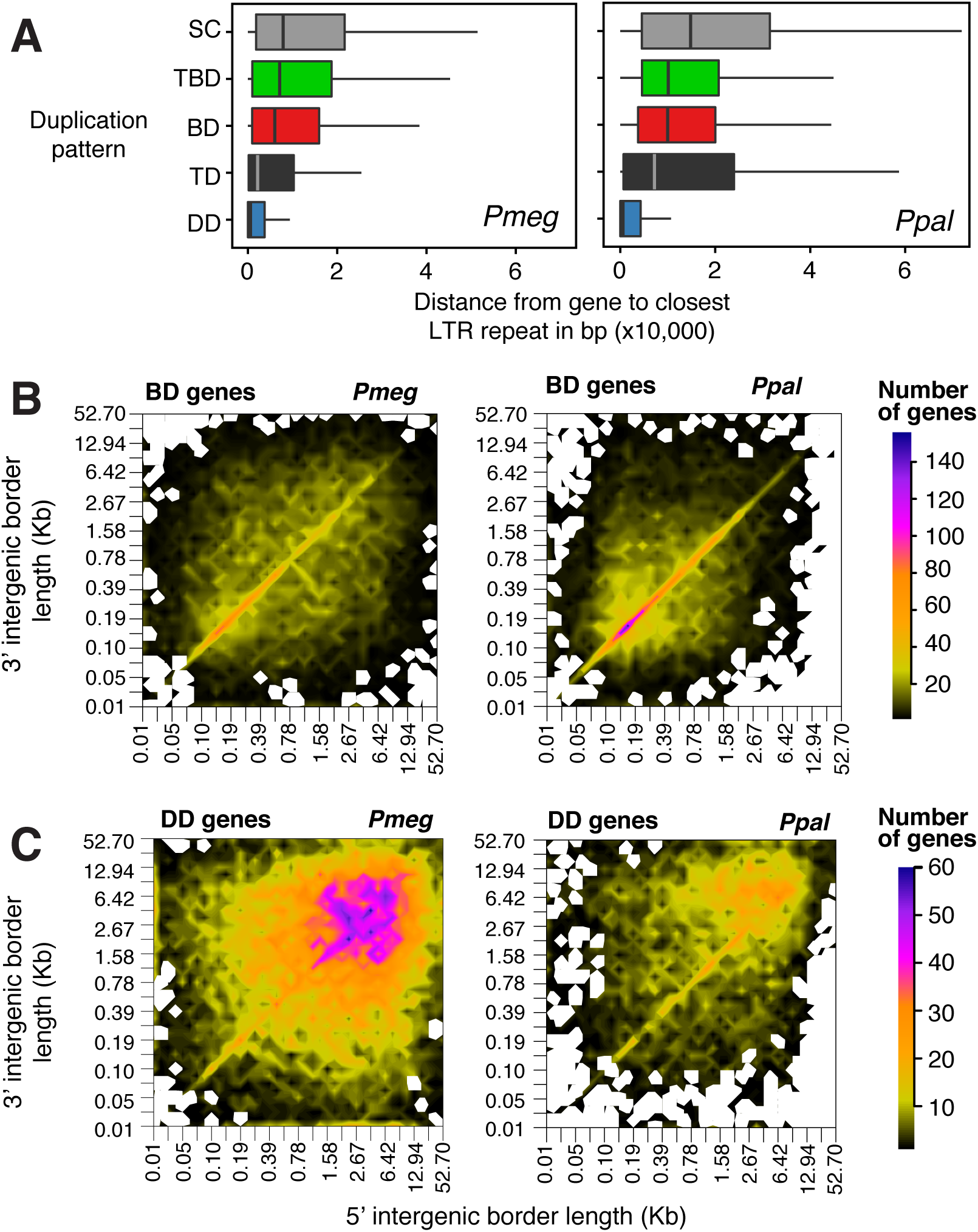
(A) Distribution of the distance between the duplication patterns and LTR TEs. (B) Intergenic space heatmap of genes in block duplicated genes (C) and dispersed duplicated genes.

On average, we found 7.4 LTR elements within 10 kbp (up or downstream) from the DD genes compared to only 4.5 elements from the BD genes (SI Appendix, Table S8). The LTRs were larger, on average, around DD (991.7 bp) and TD (949.1 bp) genes than when they were near other gene duplication categories (833.0 bp). As expected, the differential accumulation of repeats across the duplication patterns had an important impact on the intergenic space. The highest intergenic frequency of the BD genes was around 200 bp in both species (**Fig. 3B**), while the intergenic space of the DD genes had the highest frequency around 4,500 bp. DD genes had an overall trend towards the most expanded intergenic regions of the genome in both species, but it was much more striking in *Pmeg* (**Fig. 3C**).

We then inspected the association between TEs and gene families unique to or dominated by *Pmeg* and described their duplication patterns (Dataset S3). We found gene family duplicates in repetitive regions that were very close to one another. For example, 69.4% of FA00226 family members were on scaffold 20 and all 53 members of family FA00514 were on scaffold 440. There were also instances of multiple highly expanded gene families, mainly non-expressed and hypothetical protein families, loosely grouping together throughout the dispersed regions of the genome. These families were frequently associated with and overlapped LTR/Gypsy TEs (e.g.: 1,120 members of FA00003 and 877 members of FA00005), but many other consistent associations were observed as well. This included associations between FA00460 and DNA/PiggyBac and between FA00326, FA00947, and Helitrons.

We observed another gene duplication pattern, so far unique to *Pmeg*, which included multiple unique expanded gene families arranged in duplicated inverted blocks (e.g.: FA00074, FA00107, FA00211, and FA00254 and others; Dataset S3). These families primarily consist of hypothetical proteins and are associated with DNA/PiggyBac-like TEs, and often include inverted repeats on the gene block ends and repeat sequences in-between (SI Appendix, Fig. S5). DNA/PiggyBac TEs create double stranded DNA breaks, leaving TTAA overhangs not requiring DNA synthesis for repair (22). Related elements are found in diverse species (23), Inverted repeats are expected to cause instability and are possibly associated with the required chromosomal rearrangements needed to stabilize the genome after WGD.

Overall, when looking at many highly expanded gene families in *Pmeg* and *Ppal*, we observed a high number of hypothetical protein-coding genes, a tendency to be non-expressed, and their often-close association with TEs. On the other hand, these associations are less clear for expanded functional gene families like RxLRs in these genomes. Duplications of functional gene families have long been loosely associated with TE-driven genome expansion (24, 25), potentially carried along in part or entirely to new location in the genome, and subject to potential modification in the process. This would appear to be a major difference among the expanded gene families.

### Genome duplication and expansion lead to exceptional effector content in *P. megakarya*

Overall, both species were predicted to have exceptionally large number of RxLR effectors; a total of 1,382 were predicted in *Pmeg*, 717 in *Ppal* (**Fig. 4A**). Although the largest RxLR families were expanded in both pathogens, this was clearer in *Pmeg* than in *Ppal* (**Fig. 4B**). In addition, there were many “putative effectors” sharing homology with and often bordering RxLRs in both species (561 in Pmeg and 251 in Ppal). These putative effectors, though incomplete, are candidates for evolving or degenerating RxLRs, as is to be expected considering the expanded RxLR numbers in these two species.

**Fig. 4.**
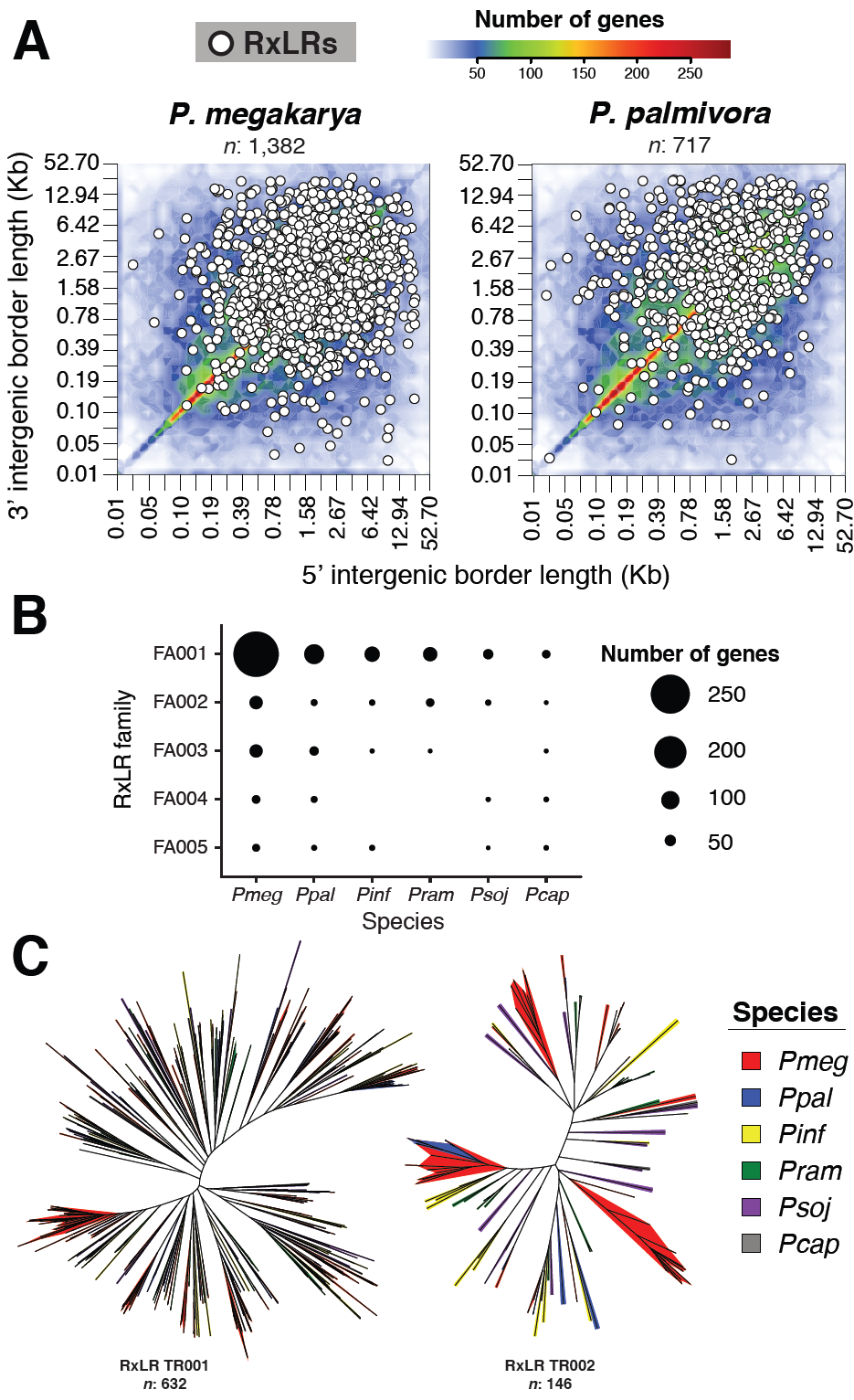
(A) Intergenic space heatmap of *Pmeg* and *Ppal* with an overlayed relative position of the predicted RxLR effectors. (B) Numbers of genes in the top five RxLR families per species. (C) Phylogenetic analysis created by resampling the site likelihoods 1,000 times of the two largest families across multiple species of *Phytophthora* showing a clear branch-specific amplification pattern in *Pmeg.*

A phylogenetic analysis of the two largest RxLR families in multiple *Phytophthora* species showed a pattern of branch-specific duplications within each gene family and the most dramatic expansions in *Pmeg* (**Fig. 4C**). The most expanded RxLR gene family (RxLR-FA001) accounted for 21.4% and 15.6% of the RxLRs in *Pmeg* and *Ppal*, respectively. Many of the RxLR effectors shown in Figure 4C were found in the expanded gene-sparse regions of the genome in both pathogens. However, in contrast to the pattern of intergenic space surrounding dispersed duplicated genes in general (**Fig. 3C**), many RxLR effectors were also abundant in the more compact (interegenic space ≤ 10 kbp) and gene-rich regions of the genome (**Fig. 4A**). This suggests that both species have evolved large effector repertoires by both WGD and expansion.

Other gene families with virulence related functions were also detected among the expanded families. Necrosis inducing proteins (NPP1, FA00029) and a family of bZIP transcription factors (FA00017) required for zoospore motility and plant infection (26, 27), are known to be expanded in *P. sojae* (28), and were also within the top 10 most-expanded gene families in both species. In *Pmeg*, a serine/threonine protein kinases family (FA00009) expanded to a size 169 genes larger than other species. These kinases are important for zoospore release, zoospore viability, encystment, and cyst germination and could be related to the higher sporulation ability of *Pmeg* (29).

Genes within RxLR families that were unique to or expanded in *Pmeg* (FA047, FA107, FA147) or *Ppal* (FA033, FA153, FA179) were induced *in planta* or in zoospores. At least 1,058 of 1,381 *Pmeg* Pm1 RxLRs and 540 of 717 *Ppal* Pp2 RxLRs were transcriptionally active (Dataset 1, SI Appendix, Table S9 and Fig. S6-S8). Twenty-two RxLRs in *Pmeg* and 32 RxLRs in *Ppal* were consistently detected as up-regulated *in planta* compared to their expression in mycelia or zoospores (SI Appendix, Fig. S9).

Together, these data show that genome duplication and gene family expansion led to increase of genes with functions related to virulence, potentially enabling these pathogens to adapt to novel circumstances.

### Genetic signatures point to adaptive evolution in the expanded gene families

Genetic analyses of multiple individuals are necessary to evaluate the evolutionary effects of WGD and gene family expansion in *Ppal* and *Pmeg*. Fifteen isolates of *Pmeg* from three West African countries and 18 isolates of *Ppal* from diverse sources were sequenced (SI Appendix, Table S1 and Fig. S10). Next, synonymous (*d*_S_) and nonsynonymous (*d*_N_) substitutions rates were estimated in pairwise comparisons of individuals for genes potentially under positive selection; such genes had an Omega ratio (ω *= d*_N_*/d*_S_) greater than one. In total, 11,716 *Pmeg* Pm1 genes and 6,567 *Ppal* Pp2 genes had ω > 1.

We then tested for evolutionary signal enrichment among > 2-fold expanded families with the Bonferroni-corrected Fisher’s Exact Test (BCFET). Remarkably, genes under positive selection in both cacao pathogens were significantly enriched in the largest RxLR gene family (“FA00008”, BCFET *P* < 2.01 × 10^−10^), an expanded bZIP transcription factor family (“FA00017”, BCFET *P* < 1.11 × 10^−9^), and an expanded M96 mating-specific family (“FA00043”, BCFET *P* < 1.30 × 10^−2^). The significant enrichments of positively selected genes in specific highly expanded families strongly suggests that their expansion is evolutionarily beneficial.

Next, SNPs in the population associated with premature stop codons were screened to identify genes likely under relaxed selection. We found 10,702 and 5,289 genes with a gained stop codon in *Pmeg* Pm1 and *Ppal* Pp2, respectively. The RxLR-encoding family (“FA0008”) was enriched in genes potentially under relaxed selection in both species (BCFET *P* < 1.73 × 10^−2^). Among the families enriched in genes potentially under relaxed selection, we also found an additional expanded RxLR family in *Pmeg* (“FA00137”, BCFET *P* < 1.30 × 10^−2^) and an expanded CRN effector family in *Ppal* (“FA00255”, BCFET *P* < 3.34 × 10^−2^). However, most (∼80%) of the families significantly enriched with premature gained stopped codons (BCFET *P* < 0.05) and potentially under relaxed selection were designated “hypothetical protein”. The large number of genes in expanded families with predicted premature gained stop codons suggests that relaxed selection is permitted by large-scale duplicative processes that increase redundancy.

## DISCUSSION

The genomes of *Pmeg* and *Ppal* revealed information about their evolution and significant differences between the two pathogens that might explain the high virulence of *Pmeg* in cacao compared to *Ppal*. Our analyses support the hypothesis that *Pmeg*, like *Ppal* (4), also underwent WGD. Evidence of WGD includes (*i*) genomes larger than other *Phytophthora* species, which was confirmed for multiple isolates and validated with flow cytometry, (*ii*) a high number of genes duplicated in conserved colinear blocks, (*iii*) a large number of expanded gene families, and (*iv*) the distribution of Ks mutations around a single event. Together, the flow cytometry results and genome structure confirmed that both species are diploid and that WGD was followed by diploidization for both species (i.e.: paleopolyploids)(30). The differences in the number of chromosomes between *Pmeg* (5-6 chromosomes) and *Ppal* (9-12 chromosomes) may have to do with the re-diploidization process after the WGD event (15, 16).

A WGD event can occur as autopolyploidy by doubling the copy number of each chromosome, or as allopolyploidy by the hybridization of two different species (30, 31). Both allo- and autopolyploidization have been reported in *Phytophthora* (32, 33) and an ancient WGD event has been suggested for some *Phytophthora* species (34). However, inter-species hybridization of *Pmeg* or *Ppal* have not been observed. The single peak in the Ks distributions in both species does not suggest that genome duplication was caused by the hybridization of distinct species, but rather that WGD was due to autopolyploidization. Our estimates suggest that the WGD event happened independently and much later than the divergence between the species, with the *Ppal* duplication occurring ∼0.65 MYA before *Pmeg*. The timing of the WGD events provides an opportunity to study how recent WGD may influence pathogen virulence. Polyploid isolates of *Pinf* (31) and hybrid *P. alni* subsp. *Alni* (32) are reportedly better adapted to specific environments than diploid *Phytophthora*. In plants, the incidence of polyploidy is greater in populations subjected to environmental challenges like high altitudes and large-scale climatic perturbation (33). The ability of the polyploids to occupy new habitats is well-documented in plants and fishes (34–36); while in plant pathogens, it has been associated with the expansion of their host range (37). WGD and the subsequent modification of copied sequences can enhance genetic diversity and be advantageous if accompanied by neo- and sub-functionalization (38–42). For example, in maize nearly 13% of duplicated genes after a WGD have functionally divergent regulatory regions (i.e. neo-functionalization) and are expressed at novel times and environments (41), while an ancient WGD in ray-finned fishes is reportedly responsible for over 10% of the biodiversity in this group of fishes (35).

In *Pmeg* and *Ppal*, we show that much larger families of virulence factor genes (4) compared to other *Phytophthora* species (7, 8, 43, 44), including RxLR, CRN and NPP1 effectors, are the result of WGD and gene expansion. Remarkably, the process of WGD and gene family expansion makes *Pmeg* the *Phytophthora* species with the highest number of RxLRs effectors observed so far. Effectors facilitate host colonization by modulating the plant immunity (45). Effector genes often undergo rapid changes in pathogen populations to overcome newly-evolved host resistance molecular mechanisms, thus the rapid effector diversification is a crucial component of pathogen success (46, 47). Our population genetic analysis confirms that the largest families of RxLRs and other expanded families in both species were overrepresented among genes under positive selection. This suggests that these genes and the gene family expansion participate in the adaptive evolution of the pathogens by creating new beneficial traits in the population (48, 49). Similar results have been observed in *P. sojae* and *P. ramorum*, in which the C-terminal effector domain of the RxLRs proteins are under positive selection (50). Genes under positive selection can confer essential traits, like the ability to escape host recognition through the novel versions of the effector protein, as shown in the flax rust fungus (*Melampsora lini*) (51). In the *Phytophthora* genus, potential disease-related genes like the phytotoxin-like scr74 gene family in *Pinf* (52) and effectors from the CRN family in *Psoj* (53) have also been reported to be under positive selection.

Although we found evidence for WGD duplication events for both species, we show that *Pmeg* has a larger genome, a greater number of genes, and a greater number of duplicated genes that are found dispersed in the genome. Dispersed duplicated genes account for most of the difference between the number of genes and genome sizes of *Pmeg* and *Ppal*. Most dispersed duplicates were tightly associated with transposable elements and tended to occur in genomic regions with the largest intergenic space. Genes with virulence-related functions were also often located in gene-sparse regions of the genome. The manner in which the dispersed duplicates are evolving is consistent with the two-speed genome mode of evolution described in other eukaryotic pathogens (54, 55), in which the gene-sparse and repeat-rich regions of the genome are the largest sources of novel functions. However, despite the great number of dispersed duplicated genes and a likely contribution of TEs in their dispersion, the intergenic space of *Pmeg* is significantly smaller than that of *Pinf*. Thus, the genome of *Pmeg* exhibits a unique architecture with a size of 222 Mbp and intergenic space smaller than 20 kbp. The unique combination of WGD and large scale transposable element driven gene/genome expansion has given *Pmeg* one of the largest genomes, the largest transcriptome, and candidate effector pools among *Phytophthora* species known to date. The combination of a genome size twice that of *Ppal* and a chromosome number half that found in *Ppal* manifests itself in the extra-large chromosomes, which is key to *Pmeg’s* identification and recognized in the species name “megakarya” so many years ago (15, 16).

This study constitutes a significant step towards unraveling the virulence differences in consequential *Phytophthora* pathogens, *Pmeg* and *Ppal.* The screening of isolates based on the effector repertoire discussed here can inform breeding programs and monitor pathogen evolution and epidemics. Additional research is required to determine the molecular causes of differences in virulence between these species and the basis of *Pmeg* aggression. The resources produced by this study should be helpful in the pursuit of this goal.

## Data availability

The sequence reported in this paper has been deposited in the National Center for Biotechnology Information Sequence Read Archive (NCBI SRA) database (accession no. PRJNA578180).

## ACKNOWLEDGMENTS

This work was supported by the Mars-USDA TRUST FUND COOPERATIVE AGREEMENT No. 58-6038-6-004 and the Mars-UC Davis agreement A18-1698. We are thankful to David Guest, Ishmael Amoako-Attah, Andrew Yaw Akrofi, B.A. Didier Begoude, G. Martijn ten Hoopen, Klotioloma Coulibaly, Boubacar Ismael Kebe, Asman Asman and J. Saul Maora for the *Phytophthora* collection. We are grateful for the comments of Brandon Gaut, Amanda Vondras, and MacKenzie Patton. References to a company and/or product by the USDA are only for the purposes of information and do not imply approval or recommendation of the product to the exclusion of others that may also be suitable.

## MATERIALS AND METHODS

### Biological material

*Pmeg* and *Ppal* isolates were obtained from collections held by *USDA, Beltsville, USA* and *Sydney Institute of Agriculture, Australia*. Isolates were initially collected from black pod infected cacao in Western Africa, Central America, Southeast Asia and Papua New Guinea and identified to species as previously described (3). Isolates of *Phytophthora* spp. and their sources are listed in SI Appendix, Table S1. The procedure for nuclei size estimates by flow cytometry were carried out as in (56), using *Arabidopsis thaliana* Col-0 as a control. Details of DNA and RNA extraction, and library preparation details are described in the SI Appendix.

### Genome assembly and annotations

SMRT reads generated on the PacBio Sequel machine were assembled using FALCON-Unzip (57), adopting the custom pipeline published in (58) and available at: https://github.com/andreaminio/FalconUnzip-DClab. Repetitive content was predicted using a combination of RepetModeler (59) and RepeatMasker (60). Protein-coding genes were predicted and polished with BRAKER (61) and PASA (62) pipelines using assembled transcriptomes as evidence. HMM models generated from classified gene families from RxLR effectors of published *Phytophthora* genomes (**Table 2**), and the HHBlits iterative sequence searcher (63) were used to predict RxLR effectors. For more detail information about the genome assembly and annotations see SI appendix.

### Duplication pattern analyses

Paralogs and orthologs were predicted using Orthofinder (64) across all the *Phytophthora* species studied. MCScanX (65) was used to classify duplicated genes into each intra-species duplication patterns. The program KaKs_Calculator v2.0 (66) was used to create synonymous mutation (Ks) profiles. For more detail information about the duplication pattern analyses see SI appendix.

