## Supplementary material for "The unique genome architecture of the devastating West African cacao black pod pathogen *Phytophthora megakarya*": S1 Appendix

**This PDF file includes:**

Supplementary text
Figures S1 to S10
Tables S1 to S9
Legends for Datasets S1 to S3
SI References

#### SUPPLEMENTARY INFORMATION TEXT

##### Supplementary Methods:

**Isolation of *Pmeg and Ppal* gDNA for short-read sequencing.** For DNA extraction, *Pmeg and Ppal* isolates were grown for seven days on 20% clarified V8 agar (CV8). 2-3 agar plugs (0.25 $cm^2$) from the cultures were then transferred to 50 ml falcon tubes containing 20 ml liquid CV8 and grown at room temperature while shaking at 100 rpm. Cultures were incubated for 5-10 days. The mycelia were washed with sterile water and collected by centrifuging at 20,000 g for 10 min, followed by flash freezing in liquid nitrogen and freeze dried. Approximately 0.1 g of freeze-dried mycelia were pulverized in a mortar and pestle under liquid nitrogen. DNA was extracted by adding the ground mycelia to a tube containing 10 ml of pre-warmed (65°C) modified CTAB buffer (3% cetyltrimethylammonium bromide, 100 mM Tris, pH 8, 20 mM EDTA, pH 8, 1.4 M NaCl, 1% PVP 40,000, 0.2% 2-mercaptoethanol, 80 µg $ml^{-1}$ proteinase K) followed by incubation at 65°C for 1 h. Each sample was extracted twice with 10 ml chloroform and the upper phase was transferred to a fresh tube. DNA was precipitated by adding 5 ml 7.5M ammonium acetate and 20 ml absolute ethanol to each tube and holding on ice for 60 min followed by centrifuging at 18,000 g for 15 min. The DNA pellet was washed with 70% ethanol, air-dried and re-suspended in 500 µl EB buffer (QIAGEN, USA). For RNase treatment, samples were treated with 0.5 µg/µl RNase A (Invitrogen, USA) enzyme followed by incubation at 37°C for 15 min. Samples were again subjected to chloroform extraction and ethanol precipitation as mentioned above and DNA was re-suspended in 100 µl EB buffer.

**Isolation of *Pmeg and Ppal* gDNA for SMRT sequencing.** High molecular weight gDNA was extracted using a modified protocol described by Stoffel *et al.* (1). Freeze dried mycelia (≃ 0.25 g) were grind in a mortar and pestle under liquid nitrogen and transferred to 25 ml chilled stainless-steel jar. The jar was submerged in liquid nitrogen for few minutes and transfer to TissueLyser II (Qiagen, USA). Sample was further grind for 45 s at 30 Hz. Grind tissue were transfer to 50 ml falcon tube containing 15 ml of pre-warmed 2X extraction buffer (100 mM Tris–HCl pH 8.0, 1.4 M NaCl, 20 mM EDTA, 2% w/v CTAB, 10 µl/ml β-mercaptoethanol), gently mixed and incubated at 65°C for 1 h. The tube was centrifuged at 10,000 rpm for 10 min to remove the cellular debris and the supernatant was transferred to a new tube containing 15 ml chloroform:isoamyl alcohol (24:1) (ChlA), gently mixed, and centrifuged at 10,000 rpm for 30 min. The aqueous phase was then transferred to a new tube containing 7.5 ml of 5 M NaCl and equal volume of ChlA was added and mixed gently and centrifuged at 10,000 rpm for 10 min. The aqueous phase was then transferred to 3 Oak Ridge tubes and 4 to 5 volumes of precipitation buffer (50 mM Tris–HCl pH 8.0, 10 mM EDTA, 1% w/v CTAB) were added. The sample was incubated overnight at room temperature to precipitate the DNA and then centrifuged at 14,000 rpm for 30 min. The DNA pellet was washed with 5 ml $dH_2O$ and pulled in to one Oak Ridge tube and centrifuged at 14,000 rpm for 10 min. DNA pellet was dissolved in 500 µl of 1.5 M NaCl and 1 µg/µl RNaseA (Invitrogen, USA) was added to the pellet and incubated at 37°C for 45 min. A chloroform extraction was performed as above to remove RNaseA and any additional contaminants. The aqueous phase was collected, and DNA was precipitated with 3 volume of absolute ethanol, followed by centrifugation for 30 min at 14,000 rpm and washed with 70% ethanol. The air-dried pellet was re-suspended in 100 µl EB buffer (QIAGEN, USA). The sample was diluted 1:25 and concentration of the gDNA was quantified with a Qubit 3.0 fluorometer using a Qubit dsDNA HS Assay Kit (Thermo Fisher Scientific, USA). The quality of the extracted gDNA was assessed using a NanoDrop UV/Vis spectrophotometer and 1% (w/v) agarose gel. Approximately 1 µg of the gDNA was run on a 0.75% pippin pulse (Sage Science, USA) gel to examine the integrity and molecular weight of the gDNA.

**Isolation of RNA from mycelia and zoospores.** For RNA extraction from mycelia (*Pmeg* and *Ppal* isolates Pm1 and Pp2 respectively), 2-3 agar plugs from a V8 agar plate culture were transferred to 250 ml conical flasks containing 50 ml liquid CV8. Liquid cultures were grown 7 days at room temperature (≃25°C) with shaking at 100 rpm. Mycelia were washed with sterile water and collected by centrifuging at 20,000 g for 10 min followed by flash freezing in liquid nitrogen and freeze drying. Freeze-dried mycelia were ground in a mortar and pestle in liquid nitrogen and transferred to a 50 mL centrifuge tube containing 15 mL of 65°C extraction buffer (2). The remaining extraction procedure was conducted as described (3). Using a NanoDrop spectrophotometer (Thermo Scientific, USA), RNA concentrations were determined based on absorbance at 260 nm and purity was estimated by the 260/280 and the 260/230 ratios.

For RNA extraction from zoospores, cultures of Pm1 and Pp2 were grown on a CV8 agar plate for 7 days under constant darkness at 25°C and then transferred to constant light (200 lux) for 4-5 days. For zoospore release, each plate was flooded with cold sterile water (4°C) and kept at 4°C for 45 min then transferred to 28°C for 28 min. The zoospore suspension was transferred to sterile 50 ml falcon tubes and collected by centrifuging at 15,000 g for 5 min. Zoospore pellets were transferred to a mortar and pestle and ground in liquid nitrogen. RNA extraction was carried out as described for mycelia.

**RNA extraction from infected plant material.** Harvested pods of the susceptible cacao clone 'Catongo' were cut into 2.5 X 2.5 cm pieces; the inner core materials were removed and the pieces were surface-sterilized with 6% ($vv^{-1}$) bleach (Clorox, USA) for 90 s followed by three rinses with sterile distilled water. Pieces were placed in sterile plastic containers (20 X 10 X 6 cm) lined with sterile tissue paper soaked in 0.7 mM benzimidazole solution (as a senescence retardant) at the bottom. For inoculation, 1 $cm^2$ sterile Whatman no. 2 filter papers were soaked in the zoospore solutions ($10^5$ zoospores $ml^{-1}$) and placed in the middle of the exterior part of each husk piece. Control husk pieces were treated with filter paper soaked in sterile water. Containers were covered and incubated at 25°C with 50% relative humidity under 12 h light (200 lx) and dark cycles. At 15 and 36 h post inoculation, the Whatman filter papers were removed and the husk pieces flash frozen in liquid nitrogen followed by freeze drying. For RNA isolation, freeze dried material was ground finely and approximately 0.05 g material transferred to a 50 mL centrifuge tube containing 15 mL of 65°C extraction buffer and RNA extraction was carried out as described for mycelia.

**Whole genome and transcriptome libraries preparation.** For preparation of DNAseq libraries, Bead-cleaned genomic DNA was randomly sheared to around 450 bp with a Covaris E220 sonicator. Sheared DNA was end-repaired, A-tailed and ligated to single adapters using the Kapa LTP library prep kit (Kapa Biosystems). RNAseq libraries were prepared using the Illumina TruSeq RNA sample preparation kit v.2 (Illumina, CA, USA), following Illumina's protocol (Low-throughput protocol) and barcoded individually. Final libraries were evaluated for quantity and quality with the High Sensitivity chip in a Bioanalyzer 2100 (Agilent Technologies, CA) and Qubit (Invitrogen, CA). DNA libraries were submitted for sequencing at 150-bp paired-end mode on an Illumina HiSeq4000 sequencer (Novogene Co. Ltd., Beijing, China).

**SMRTbell Libraries Preparation.** When needed, 200 µl of HMW gDNA at a concentration of 100 ng/µl were fragmented using a 26G blunt needle (SAI Infusion Technologies). gDNA shearing was done by aspirating the entire volume and passing the sample through the 26G blunt needle fifteen times. After shearing, sample was cleaned and concentrated using 0.45X AMPure PB beads and size distribution of the sheared gDNA fragments was evaluated using pulse field gel electrophoresis (Pippin pulse, Sage Science) prior to libraries preparation. SMRTbell template libraries from Pm1 and Pp2 isolates were prepared with 6-12 µg of sheared DNA using SMRTbell Template Prep Kit (Pacific Biosciences) following the manufacturer's instructions. 30 ul of SMRTbell template was loaded into the Sage Blue Pippin for size selection and a cutoff range of 17-50 Kbp was selected. Size selected library was cleaned with 1X AMPure PB beads and a repair DNA damage treatment was performed. After DNA repair, a new clean up step with 1X AMPure PB beads was done. A total of 6 SMRT cells per isolate were sequenced on the PacBio sequel system (Novogene Co. Ltd., Beijing, China). SMRTbell template libraries from Pm4, Pm15, Pp3, and Pp15 isolates were prepared with 6 µg of sheared DNA using the SMRTbell Express Template Prep Kit v2.0 (Pacific Biosciences) following the manufacturer's instructions. A cutoff range of 17-50 Kbp was chosen for size selection. Size selected library was cleaned with 1X AMPure PB beads. 4 SMRT cells per isolate were sequenced on the PacBio sequel system using a V3 chemistry (Genome center, UCDavis).

**Genome assembly.** Assembly of the genomes was performed using SMRT reads with FALCON-Unzip ver. 2017.06.28-18.01 (4) adopting the custom pipeline published in (5). The pipeline code is available at https://github.com/andreaminio/FalconUnzip-DClab. Before performing error correction of the raw reads, repetitive regions were marked using the TANmask and REPmask modules from the DAmasker v1.0 (6), reducing the complexity of the read-to-read alignment phase. After error-correction, reads were again marked before proceeding with the assembling phase. This additional repeat masking step increased assembly contiguity by reducing the complexity of the overlap graph. FALCON was performed using different thresholds on seed-reads minimum length for overlap stage (length_cutoff_pr parameter) in order to find the least fragmented primary assembly. Parameter set include: "falcon_sense_skip_contained = TRUE", "falcon_sense_option = --output_multi --min_idt 0.70 --min_cov 4 --max_n_read 400", "length_cutoff_pr = 27000", "ovlp_DBsplit_option = -x500", "ovlp_HPCdaligner_option = -mtan -mrep2 -v -B128 -M60 -t60 -k20 -h256 -e.9 -l1000 -s100 -T16", and "overlap_filtering_setting = --max_diff 100 --max_cov 400 --min_cov 3". Unzip procedure for haplotype phasing was carried out with default parameters (4), followed by polishing of primary contigs and haplotigs with Arrow (from ConsensusCore2 v.3.0.0) using long reads. Primary contigs were scaffolded using SSPACE-Longreads v.1.1 (7), followed by gap closing with PBJelly (PBsuite v15.8.4; (8, 9)). To assess the genome assembly length, SMRT reads of *Pmeg* isolate Pm1 and *Ppal* isolate Pp2 were also assembled using two other alternative assemblers, Canu v1.8-14 (10)) and WTDBG2 v2.3 (11). Canu was performed separately by setting an expected genome size of 215 Mbp and 115 Mbp, error correction, trimming and assembly parameters are: "Error correction: minReadLength=1000 minOverlapLength=500 corOutCoverage=80", "Trimming: minReadLength=1000 minOverlapLength=500", and "Assembly: correctedErrorRate=0.04 minReadLength=1000 minOverlapLength=500". WTDBG2 was performed with the parameters include "-S 4 -p 21 -k 0 -e 4 -L 5000" and consensus called with wtpoa-cns algorithm. Assembled sequences were then polished and scaffolded using using long reads with Arrow (from ConsensusCore2 v.3.0.0) and SSPACE-Longreads v.1.1 respectively. The gene space completeness of the genome assemblies was evaluated using the BUSCO genes (12), specifically using BUSCO v1.22 and the "eukaryota_odb9" database.

**Nuclei size content inference by flow cytometry.** The details of the procedure can be found elsewhere (13). In brief, stationary grown mycelia in test tubes containing 5 mL filter-sterilized 5% clarified V8 broth for 7 days at 25°C were harvested. Approximately 1 mg of dry blotted sample and five flower buds of *Arabidopsis thaliana* Col-0 were then combined and co-chopped in a Petri dish containing 500 µL extraction buffer (Cystain PI absolute P Kit, Sysmex America Inc.), and the suspension was filtered through a 10 µm filter (CellTrics, Sysmex America Inc.) and 2 mL of Propidium Iodine staining solution was added (14, 15). Measurements were done on a Becton Dickinson FACScan (Franklin Lakes, NewJersey) equipped with a 488 nm laser and a 585/42 nm band pass filter. Three biological replicates were measured per isolate. The data were analyzed using FlowJo v.10 (https://www.flowjo.com/solutions/flowjo) and DNA content was inferred using a linear regression with the ratios between the peak positions of *Phytophthora* sample and the *Arabidopsis* size standard (1C = 157 MB) (14).

**Repeat prediction.** An initial run of RepeatMasker v4.0.6 (16) with the standard library was done in the new *Pmeg* and *Ppal* assemblies. The repeats predicted by this run were combined with the repeats extracted from multiple published *Phytophthora* masked assemblies (**Table 2**) to create a new database using the *de-novo* repeat predictor RepetModeler v1.0.11 (17). The classified consensus sequences were then used as a custom library for another run of RepeatMasker in the *Pmeg* and *Ppal* assemblies to predict the final repeats (Dataset 2).

**Gene prediction.** RNAseq paired-end reads of 150 bp in length and were trimmed with Trimmomatic v0.36 (18) with a 4-base wide sliding window, cutting when the average quality per base drops below 15 and a minimum length of 100 bp. Four biological replicates of each condition (i.e., mycelium, zoospores, 15 hpi or 36 hpi) were merged and assembled separately by Trinity v2.6.5 (19) with the "--normalize_reads" option to create four transcriptomes per species. The assembled transcripts were then stringently mapped with PASA v2.0.2; (20) with a minimum percentage aligned of 95% and a minimum average percentage identity of 90% using the GMAP v2015-11-20 (21) and BLAT v36x2 (22) aligners to each genome. Candidate coding regions were then searched by TransDecoder v3.0.1 (23) in the assembled and aligned transcripts within the PASA pipeline. The alignments of best scoring transcripts were used to create a high-quality set of transcripts. The splicing information of this high-quality transcripts were used as hint for the gene predictor BRAKER1 v1.9 (24) with the "--fungus" option to allow for potential overlapping genes on the softmasked genome assemblies. The resulting protein-coding genes were then taken through the PASA gene refinement pipeline, in which the initial Trinity assembled reads were mapped with more relaxed parameters (minimum alignment of 90% and minimum average identity of 85%) to polish the gene predicted models. To filter potential TEs from the predicted gene a series of filters were imposed. First, any gene that overlapped 90% or more with transposable element predicted by RepeatMasker was removed. Then, all the genes with a hit to an HMM repeat model of Dfam v3.0 (25) was also removed. Also, genes with descriptions from BLAST2GO (see below) matching "transposon”, “transposable”, “transposase”, “retrov” or “helitron"; or with PFAM domains matching Pfam "Integrase” “RVT_”, “rve”, “Retrotrans”, “gag”, ”Chromo”, “RT_”, “Helitron” or “DDE_" were filtered out. Finally, a manual filtering of genes based other functional annotations of repeats was done. Gene families were created with a reciprocal BLASTP with an e-value of 1e-10 and clustered with MCL v14-137 (26) with and inflation value of 1.5.

**Gene functional annotation.** The filtered proteins were aligned with BLASTP v2.6.0+ (27) to the whole “RefSeq” protein database. The resulting alignments were used as input into Blast2GO v4.1.7 (28) and the proteins descriptions were extracted. The predicted proteins were also searched the Pfam database v32.0 (29) using and a domain e-value cutoff of 0.001. Carbohydrate-active enzymes (CAZymes) were annotated using dbCAN2 (30) with default parameters, and using only annotations validated by two of three tools. Candidate biosynthetic gene clusters were predicted with AntiSMASH Fungal v4.2.0 (31). Proteins were classified as secreted if they had a signal secretion peptide predicted by SignalP v4.1 (32), but did not have a transmembrane domain (TM) in the first 60 amino acids or more than 2 TMs in total predicted by TMHMM v2.0 (33), and if the protein did not have a mitochondrial targeting peptide (mTP) predicted by TargetP v1.1 (34). All functional annotations can be found in Dataset S1.

**Prediction of effectors.** RxLR protein effectors from previously published *Phytophthora* (Table 2) species were classified in tribes using a reciprocal BLASTP v2.6.0+ with a cutoff e-value of 0.01, followed by a clustering with MCL using an inflation value of 1.5. The genes of each tribe with more than one member were aligned with MAFFT v7.310 (35) to create a multiple sequence that was then used as input to hmmbuild from HMMER v3.2.1 (36) to create a an HMM model per tribe. Each of these models were used to search throughout the predicted secreted proteins with a e-value of 0.01 for RxLR in the new genomes. In addition, HHBlits v3.1.0 (37) was used in combination with the clustered and deeply annotated Uniclust30 database v2018_08 (38). HHblits is the HMM-HMM-based iterative sequence search that was shown to be very sensitive in detecting threedimensional homologous proteins. Thus, the predicted secreted proteins were evaluated with HHBlits for structurally similar RxLR effectors in the new genomes compared to well annotated and validated RxLR effectors. Hits with the previous *Pmeg* or *Ppal* proteins or with RxLR-like proteins were discarded. To predict the Crinklers effectors we took previously annotated and published genes Crinklers (39–41), grouped into tribes, created HMM models per tribed and predicted the effectors based on the models in the same way than described above for the RxLRs.

**Identification of orthologs and duplicates.** The software Orthofinder v2.2.6 (42) was used to predict orthologs across the seven *Phythophtora* species. Orthofinder was used with BLASTP for the sequence search, MAFFT v7.310 (35) for the multiple alignments, FastTree v2.1.10 (43) for the tree inference method and an inflation parameter of 1.5. Within-species duplicates were defined as genes with multiple genes per paralogous groups were combined genes of the same species that had 50% or higher sequence identity, and both query and subject length coverage higher than 50% using BLASTP for the alignments. Once the duplicates were defined, a pair-wise homologous list was created of all the duplicates per species. This list was used as input for MCScanX (44) to classify the duplicates into dispersed, tandem, blocks or tandem-blocks.

**Synonymous substitution rates across duplicates**. Proteins with a within-species duplicate of 50% identity or higher were used to calculate synonymous mutations. Each duplicate sequence was aligned using MAFFT v7.310 (35) and the alignments were edited with gBlocks v0.91b (45), to remove any gaps and keep the conserved blocks. The results format gBlocks edited and used as input into KaKs_Calculator v2.0 (46). The $\gamma$-MYN algorithm was used as the method to estimate Ks using the standart genetic code table.

**Phylogenetic analysis.** Multiple sequences were aligned with MAFFT v7.310 (35) and edited by gBlocks v0.91b (45). The clock-calibrated phylogenetic done as in (47) using BEAST v1.10.4 (48). Briefly, the WAG substitution model was used, the Yule speciation process was assumed with a uniform distribution on the birthrate (0–100; initial value 0.01), while a strict clock was modeled with an exponential prior distribution (mean 1.0, initial value 0.01). The mean and 95% CI divergence time values of the strict clock estimation reported in (47) were used as priors to calibrate our phylogenetic trees. A total of 10 million generations were created by tree. The clock calibrated tree in combination with the number of genes per gene family were used to find study the evolution genes families across multiple *Phytophthora* species with the software CAFE v4.2.1 (https://github.com/hahnlab/CAFE) (49). CAFE was run with default parameters optimizing the lambda parameter (option -s) to 0.033987 with a *P*-value cutoff of 0.01 (option -p).

**Same scaffold gene block identification and validation.** Fusion events in between colinear gene blocks were selected to design primers. A maximum length of 1000 bp was used as a threshold to select the candidates for the primer design. Those sequences were extracted from the primary alignment of both species using 2000 bp upstream and downstream the fusion event. This sequences individually served as input for Primer3plus online software (50) in detection mode. For each Fusion event a total of 5 primers pairs were generated and tested *in-silico* with the program Degenerate In-Silico PCR (dispr, https://github.com/douglasgscofield/dispr). All the primers were tested against the primary assembly of the corresponding specie allowing 1 mismatch in the first 3 bases in the 5' of the primers, as well as product size range in between 200 bp and 3000 bp. The final primers sets were selected based on a specificity of the pair and the length of the product (maximum 1300 bp). The selected primers were used in a PCR with the corresponding species DNA. The reaction mix was prepared with 1X OneTaq Standard Reaction Buffer (New England Biolabs, Ipswich, Massachusetts), 1 mg/mL BSA, 0.2 mM dNTPs, 0.2 µM of each primer, 1.25 units of OneTaq DNA Polymerase (New England Biolabs, Ipswich, Massachusetts) and 1ng of DNA in a 25µL reaction, Nuclease free water was used as negative control of the reaction. The thermocycling profile (Veriti thermal cycler, Applied Biosystems) was set to an initial denaturation at 94 °C for 4 min, 31 cycles at 94 °C for 30 s, 62 °C for 30 s, and 68 °C for 1.16 min., and a final extension at 68 °C for 10 min. The amplicons were checked in an 1.5% (w/v) agarose with Apex Safe DNA Gel Stain (Apex Bioresearch Products), using 100 bp DNA ladder (New England Biolabs, Ipswich, Massachusetts). The running conditions were 80 V for 50min.

**SNPs prediction and analyses.** Paired-end reads of 150 bp were trimmed with Trimmomatic v0.36 (18) and mapped to the reference genome using BWA mem v0.7.12-r1039 (51). Duplicated mapped reads reads were removed, the remaining were tagged and used to predict SNPs with GATK's UnifiedGenotyper (52) with a minimum base score of 20 and ploidy of 2. The positions with a lower than 5 reads supporting it or higher than 1.5x the median coverage were removed for the the downstream analysis. The predicted SNPs were used as input to GATKs' "FastaAlternateReferenceMaker" v3.5-0-g36282e4 (52) to generate alternative genomes per isolate and genes were extracted using the reference coordinates. Each gene sequence per isolate was aligned with ClustalW v2.1 (53). The multiple alignment was as input to YN00 from the PAML v4.9f package (54) with the universal genetic code to caculate the Omega values per genes. The VCF files with all sites and with all isolates per species were also used to calculate the Tajimas'D using 500 bp windows and sliding 250 bp with VCF-KIT v0.1.6 (55). The VCF files were also used in SnpEff v4.3t 2017-11-24 (56) to predict the genes with a gained stop codon.

**Gene expression analysis.** The same filtered paired-end RNAseq reads used in the gene prediction were used for the gene expression analysis. The mycelia and zoospore reads were mapped to each species genome with the splice-aware mapper HISAT2 v2.0.5 (57) using the pair-end mode and with the following arguments: "-k 1 --non-deterministic". The *in planta* samples (i.e. 15 hpi and 36 hpi) were mapped to a database that included the corresponding *Phytophthora* species in combination with the *T. cacao* criollo v2 genome (58). The count of the reads was done using the gff3 files with the "summarizeOverlaps" from the R package GenomicAlignments v1.18.1 (59), to extract the reads in the predicted exons. Estimation and statistical analysis of expression level using the count data of each gene with four replicates for each library were performed using the DESeq2 package in the R statistics suite (60). For DESeq2's default normalization method, scaling factors are calculated for each lane as median of the ratio, for each gene, of its read count of its geometric mean across all lanes and apply to all read counts.

#### SUPPLEMENTAL TEXT CITATIONS


1. K. Stoffel, *et al.*, Development and application of a 6.5 million feature Affymetrix Genechip® for massively parallel discovery of single position polymorphisms in lettuce (Lactuca spp.). *BMC Genomics* (2012) https:/doi.org/10.1186/1471-2164-13-185.
2. B. A. Bailey, M. D. Strem, H. Bae, G. A. De Mayolo, M. J. Guiltinan, Gene expression in leaves of Theobroma cacao in response to mechanical wounding, ethylene, and/or methyl jasmonate. *Plant Sci.* (2005)

https:/doi.org/10.1016/j.plantsci.2005.01.002.
3. B. A. Bailey, *et al.*, Dynamic changes in pod and fungal physiology associated with the shift from biotrophy to necrotrophy during the infection of Theobroma cacao by Moniliophthora roreri. *Physiol. Mol. Plant Pathol.* (2013) https:/doi.org/10.1016/j.pmpp.2012.11.005.
4. C.-S. Chin, *et al.*, Phased diploid genome assembly with single-molecule real-time sequencing. *Nat. Methods* (2016) https:/doi.org/10.1038/nmeth.4035.
5. A. Minio, M. Massonnet, R. Figueroa-Balderas, A. Castro, D. Cantu, Diploid Genome Assembly of the Wine Grape Carménère. *G3 Genes, Genomes, Genet.* (2019) https:/doi.org/10.1534/g3.119.400030.
6. G. Myers, Efficient local alignment discovery amongst noisy long reads in *Lecture Notes in Computer Science (Including Subseries Lecture Notes in Artificial Intelligence and Lecture Notes in Bioinformatics)*, (2014) https:/doi.org/10.1007/978-3-662-44753-6_5.
7. M. Boetzer, W. Pirovano, SSPACE-LongRead: Scaffolding bacterial draft genomes using long read sequence information. *BMC Bioinformatics* (2014) https:/doi.org/10.1186/1471-2105-15-211.
8. A. C. English, *et al.*, Mind the gap: upgrading genomes with Pacific Biosciences RS long-read sequencing technology. *PLoS One* (2012) https:/doi.org/10.1371/journal.pone.0047768.
9. A. C. English, W. J. Salerno, J. G. Reid, PBHoney: Identifying genomic variants via long-read discordance and interrupted mapping. *BMC Bioinformatics* (2014) https:/doi.org/10.1186/1471-2105-15-180.
10. S. Koren, *et al.*, Canu: Scalable and accurate long-read assembly via adaptive κ-mer weighting and repeat separation. *Genome Res.* (2017) https:/doi.org/10.1101/gr.215087.116.
11. J. Ruan, H. Li, Fast and accurate long-read assembly with wtdbg2. *bioRxiv* (2019) https:/doi.org/10.1101/530972.
12. F. A. Simão, R. M. Waterhouse, P. Ioannidis, E. V Kriventseva, E. M. Zdobnov, BUSCO: assessing genome assembly and annotation completeness with single-copy orthologs. *Bioinformatics*, 1–3 (2015).
13. M. Malar C, *et al.*, Haplotype-Phased Genome Assembly of Virulent Phytophthora ramorum Isolate ND886 Facilitated by Long-Read Sequencing Reveals Effector Polymorphisms and Copy Number Variation . *Mol. Plant-Microbe Interact.* (2019) https:/doi.org/10.1094/mpmi-08-18-0222-r.
14. J. Doležel, J. Bartoš, Plant DNA flow cytometry and estimation of nuclear genome size in *Annals of Botany*, (2005) https:/doi.org/10.1093/aob/mci005.
15. L. Bertier, L. Leus, L. D'hondt, A. W. A. M. De Cock, M. Höfte, Host adaptation and speciation through hybridization and polyploidy in phytophthora. *PLoS One* (2013) https:/doi.org/10.1371/journal.pone.0085385.
16. A. Smit, R. Hubley, P. Green, RepeatMasker Open-4.0. (2015).
17. A. Smit, R. Hubley, RepeatModeler Open-1.0 (2015).
18. A. M. Bolger, M. Lohse, B. Usadel, Trimmomatic: A flexible trimmer for Illumina sequence data. *Bioinformatics* (2014) https:/doi.org/10.1093/bioinformatics/btu170.
19. M. G. Grabherr, *et al.*, Full-length transcriptome assembly from RNA-Seq data

without a reference genome. *Nat. Biotechnol.* (2011) https:/doi.org/10.1038/nbt.1883.
20. B. J. Haas, *et al.*, Improving the Arabidopsis genome annotation using maximal transcript alignment assemblies. *Nucleic Acids Res.* (2003).
21. T. D. Wu, C. K. Watanabe, GMAP: A genomic mapping and alignment program for mRNA and EST sequences. *Bioinformatics* (2005) https:/doi.org/10.1093/bioinformatics/bti310.
22. W. J. Kent, BLAT--the BLAST-like alignment tool. *Genome Res.* (2002) https:/doi.org/10.1101/gr.229202.
23. B. J. Haas, *et al.*, De novo transcript sequence reconstruction from RNA-seq using the Trinity platform for reference generation and analysis. *Nat. Protoc.* (2013) https:/doi.org/10.1038/nprot.2013.084.
24. K. J. Hoff, S. Lange, A. Lomsadze, M. Borodovsky, M. Stanke, BRAKER1: Unsupervised RNA-Seq-based genome annotation with GeneMark-ET and AUGUSTUS. *Bioinformatics* (2016) https:/doi.org/10.1093/bioinformatics/btv661.
25. R. Hubley, *et al.*, The Dfam database of repetitive DNA families. *Nucleic Acids Res.* (2016) https:/doi.org/10.1093/nar/gkv1272.
26. A. J. Enright, S. Van Dongen, C. A. Ouzounis, An efficient algorithm for large-scale detection of protein families. *Nucleic Acids Res.* (2002).
27. C. Camacho, *et al.*, BLAST+: architecture and applications. *BMC Bioinformatics* (2009) https:/doi.org/10.1186/1471-2105-10-421.
28. S. Götz, *et al.*, High-throughput functional annotation and data mining with the Blast2GO suite. *Nucleic Acids Res.* (2008) https:/doi.org/10.1093/nar/gkn176.
29. S. El-Gebali, *et al.*, The Pfam protein families database in 2019. *Nucleic Acids Res.* (2019) https:/doi.org/10.1093/nar/gky995.
30. H. Zhang, *et al.*, DbCAN2: A meta server for automated carbohydrate-active enzyme annotation. *Nucleic Acids Res.* (2018) https:/doi.org/10.1093/nar/gky418.
31. T. Weber, *et al.*, antiSMASH 4.0—improvements in chemistry prediction and gene cluster boundary identification. *Nucleic Acids Res.* (2017) https:/doi.org/10.1093/nar/gkx319.
32. T. N. Petersen, S. Brunak, G. von Heijne, H. Nielsen, SignalP 4.0: discriminating signal peptides from transmembrane regions. *Nat. Methods* (2011) https:/doi.org/10.1038/nmeth.1701.
33. A. Krogh, B. Larsson, G. Von Heijne, E. L. L. Sonnhammer, Predicting transmembrane protein topology with a hidden Markov model: Application to complete genomes. *J. Mol. Biol.* (2001) https:/doi.org/10.1006/jmbi.2000.4315.
34. O. Emanuelsson, H. Nielsen, S. Brunak, G. Von Heijne, Predicting subcellular localization of proteins based on their N-terminal amino acid sequence. *J. Mol. Biol.* (2000) https:/doi.org/10.1006/jmbi.2000.3903.
35. K. Katoh, D. M. Standley, MAFFT multiple sequence alignment software version 7: improvements in performance and usability. *Mol. Biol. Evol.* (2013) https:/doi.org/10.1093/molbev/mst010.
36. J. M. Hancock, M. J. Zvelebil, J. M. Hancock, M. J. Bishop, “HMMer” in *Dictionary of Bioinformatics and Computational Biology*, (2014) https:/doi.org/10.1002/9780471650126.dob0323.pub2.
37. M. Remmert, A. Biegert, A. Hauser, J. Söding, HHblits: Lightning-fast iterative

protein sequence searching by HMM-HMM alignment. *Nat. Methods* (2012) https:/doi.org/10.1038/nmeth.1818.

38. M. Mirdita, *et al.*, Uniclust databases of clustered and deeply annotated protein sequences and alignments. *Nucleic Acids Res.* (2017) https:/doi.org/10.1093/nar/gkw1081.
39. B. J. Haas, *et al.*, Genome sequence and analysis of the Irish potato famine pathogen Phytophthora infestans. *Nature* (2009) https:/doi.org/10.1038/nature08358.
40. K. H. Lamour, R. Stam, J. Jupe, E. Huitema, The oomycete broad-host-range pathogen Phytophthora capsici. *Mol. Plant Pathol.* (2012) https:/doi.org/10.1111/j.1364-3703.2011.00754.x.
41. S. S. Ali, *et al.*, Phytophthora megakarya and Phytophthora palmivora, Closely Related Causal Agents of Cacao Black Pod Rot, Underwent Increases in Genome Sizes and Gene Numbers by Different Mechanisms. *Genome Biol. Evol.* (2017) https:/doi.org/10.1093/gbe/evx021.
42. D. M. Emms, S. Kelly, OrthoFinder2: fast and accurate phylogenomic orthology analysis from gene sequences. *bioRxiv* (2018) https:/doi.org/10.1101/466201.
43. M. N. Price, P. S. Dehal, A. P. Arkin, FastTree 2 - Approximately maximum-likelihood trees for large alignments. *PLoS One* (2010) https:/doi.org/10.1371/journal.pone.0009490.
44. Y. Wang, *et al.*, MCScanX: A toolkit for detection and evolutionary analysis of gene synteny and collinearity. *Nucleic Acids Res.* (2012) https:/doi.org/10.1093/nar/gkr1293.
45. J. Castresana, Selection of conserved blocks from multiple alignments for their use in phylogenetic analysis. *Mol. Biol. Evol.* (2000) https:/doi.org/10.1093/oxfordjournals.molbev.a026334.
46. Z. Zhang, *et al.*, KaKs_Calculator: Calculating Ka and Ks Through Model Selection and Model Averaging. *Genomics, Proteomics Bioinforma.* (2006) https:/doi.org/10.1016/S1672-0229(07)60007-2.
47. N. H. Matari, J. E. Blair, A multilocus timescale for oomycete evolution estimated under three distinct molecular clock models. *BMC Evol. Biol.* (2014) https:/doi.org/10.1186/1471-2148-14-101.
48. A. J. Drummond, A. Rambaut, “Bayesian evolutionary analysis by sampling trees” in *Bayesian Evolutionary Analysis with BEAST*, (2015) https:/doi.org/10.1017/CBO9781139095112.007.
49. T. De Bie, N. Cristianini, J. P. Demuth, M. W. Hahn, CAFE: A computational tool for the study of gene family evolution. *Bioinformatics* (2006) https:/doi.org/10.1093/bioinformatics/btl097.
50. A. Untergasser, *et al.*, Primer3Plus, an enhanced web interface to Primer3. *Nucleic Acids Res.* (2007) https:/doi.org/10.1093/nar/gkm306.
51. H. Li, Aligning new-sequencing reads by BWA BWA : Burrows-Wheeler Aligner. *Slides* (2010).
52. A. McKenna, *et al.*, The Genome Analysis Toolkit: a MapReduce framework for analyzing next-generation DNA sequencing data. *Genome Res.* (2010) https:/doi.org/10.1101/gr.107524.110.
53. U. Clustalw, C. To, D. O. Multiple, ClustalW and ClustalX. *Options* (2003)

https:/doi.org/10.1002/0471250953.bi0203s00.
54. Z. Yang, Paml Faq. *Culture* (2005).
55. D. E. Cook, E. C. Andersen, VCF-kit: Assorted utilities for the variant call format. *Bioinformatics* (2017) https:/doi.org/10.1093/bioinformatics/btx011.
56. P. Cingolani, *et al.*, A program for annotating and predicting the effects of single nucleotide polymorphisms, SnpEff. *Fly (Austin).* (2012) https:/doi.org/10.4161/fly.19695.
57. D. Kim, B. Langmead, S. L. Salzberg, Hisat2. *Nat. Methods* (2015) https:/doi.org/10.1038/nmeth.3317.
58. X. Argout, *et al.*, The cacao Criollo genome v2.0: An improved version of the genome for genetic and functional genomic studies. *BMC Genomics* (2017) https:/doi.org/10.1186/s12864-017-4120-9.
59. M. Lawrence, *et al.*, Software for Computing and Annotating Genomic Ranges. *PLoS Comput. Biol.* (2013) https:/doi.org/10.1371/journal.pcbi.1003118.
60. S. Anders, W. Huber, Differential expression analysis for sequence count data. *Genome Biol.* (2010) https:/doi.org/10.1186/gb-2010-11-10-r106.

### SUPPLEMENTARY FIGURES

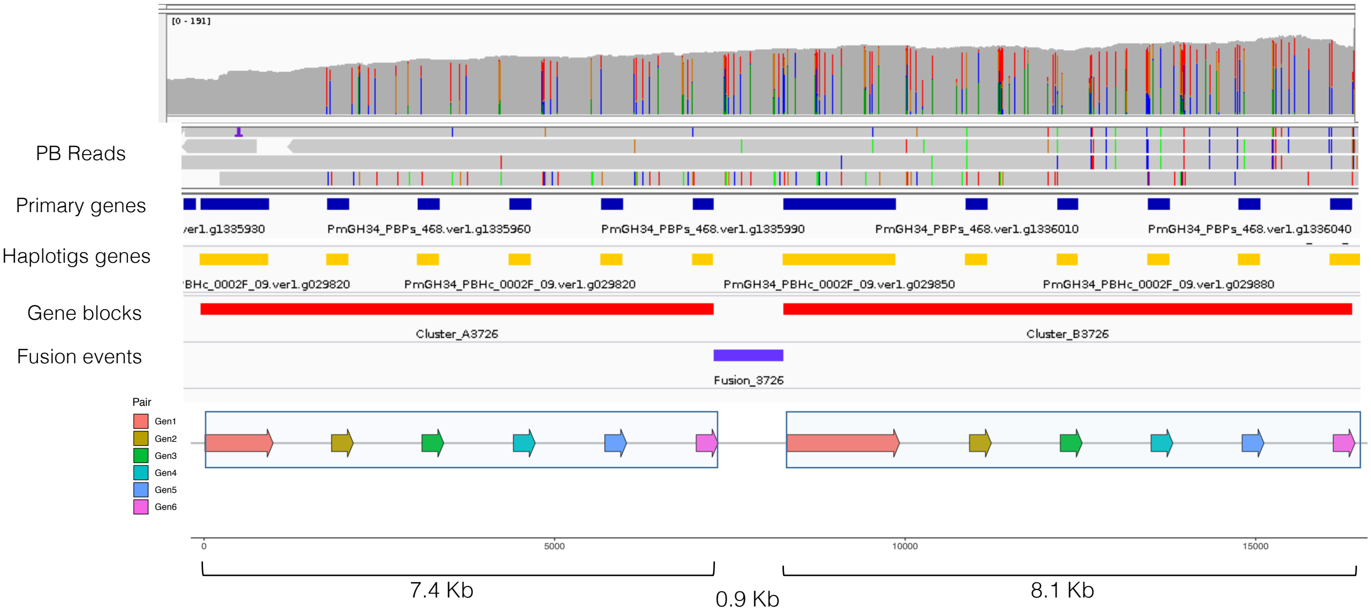


**Fig. S1.** Example of duplicated block in the same assembled primary scaffold with corresponding duplicated gene blocks in the haplotigs.

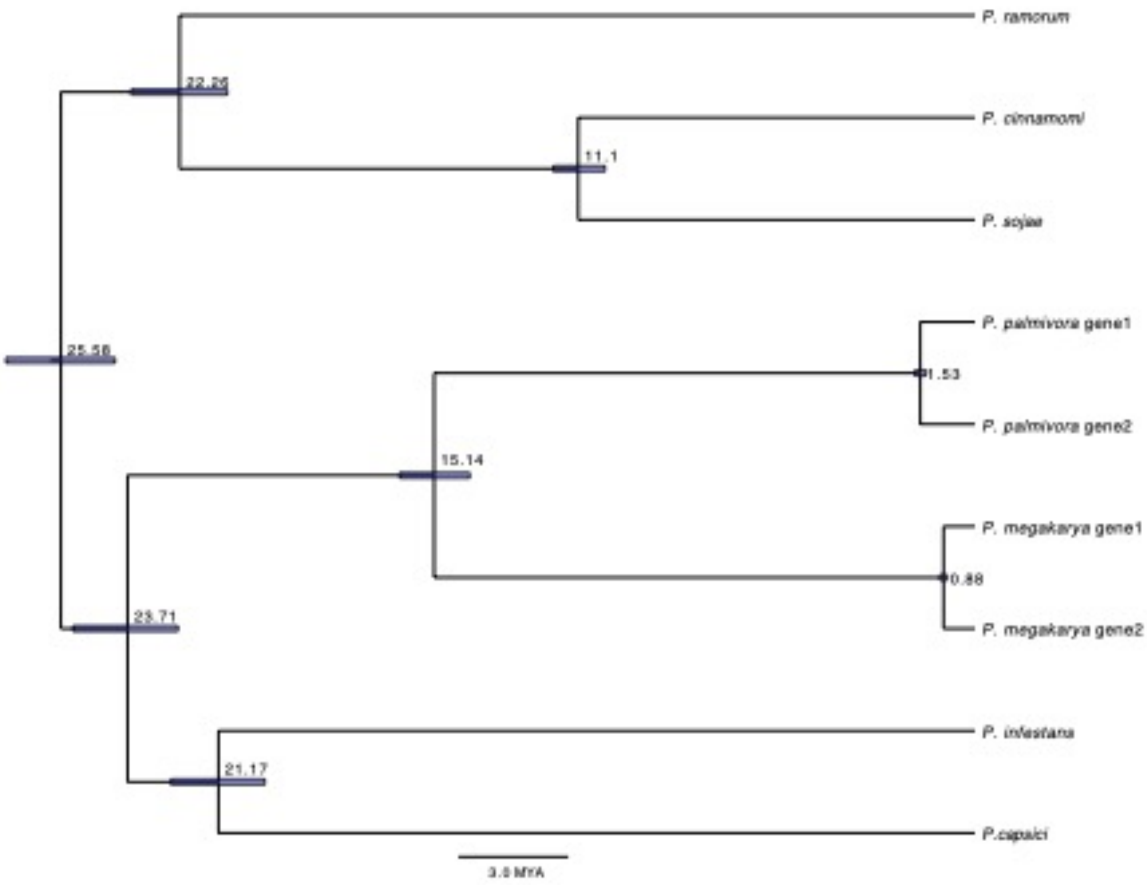


**Fig. S2.** Clock-calibrated phylogenetic tree from 215 genes detected within duplicated blocks and with exactly two copies in the genomes of *Pmeg* and *Ppal* but only one copy in the rest of the species. The presence of the duplicated genes still within the duplicated blocks strongly suggest that they were duplicated in the WGD, thus the timing of their divergence is good estimate for the WGD event.

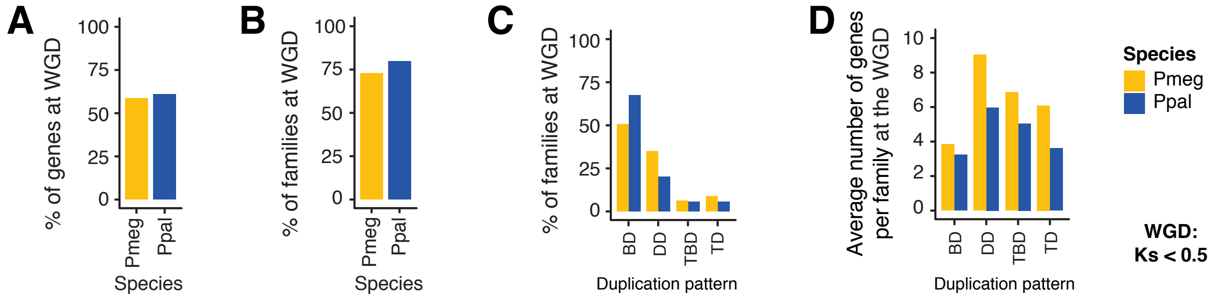


**Fig. S3.** (A) Percentage of genes estimated to have been duplicated at the WGD event (approximately Ks < 0.5). Percentage of gene families represented at the WGD event overall (B) and divided by duplication pattern (C). Average number of genes per gene family duplicated at the WGD event and separated by duplication patterns.

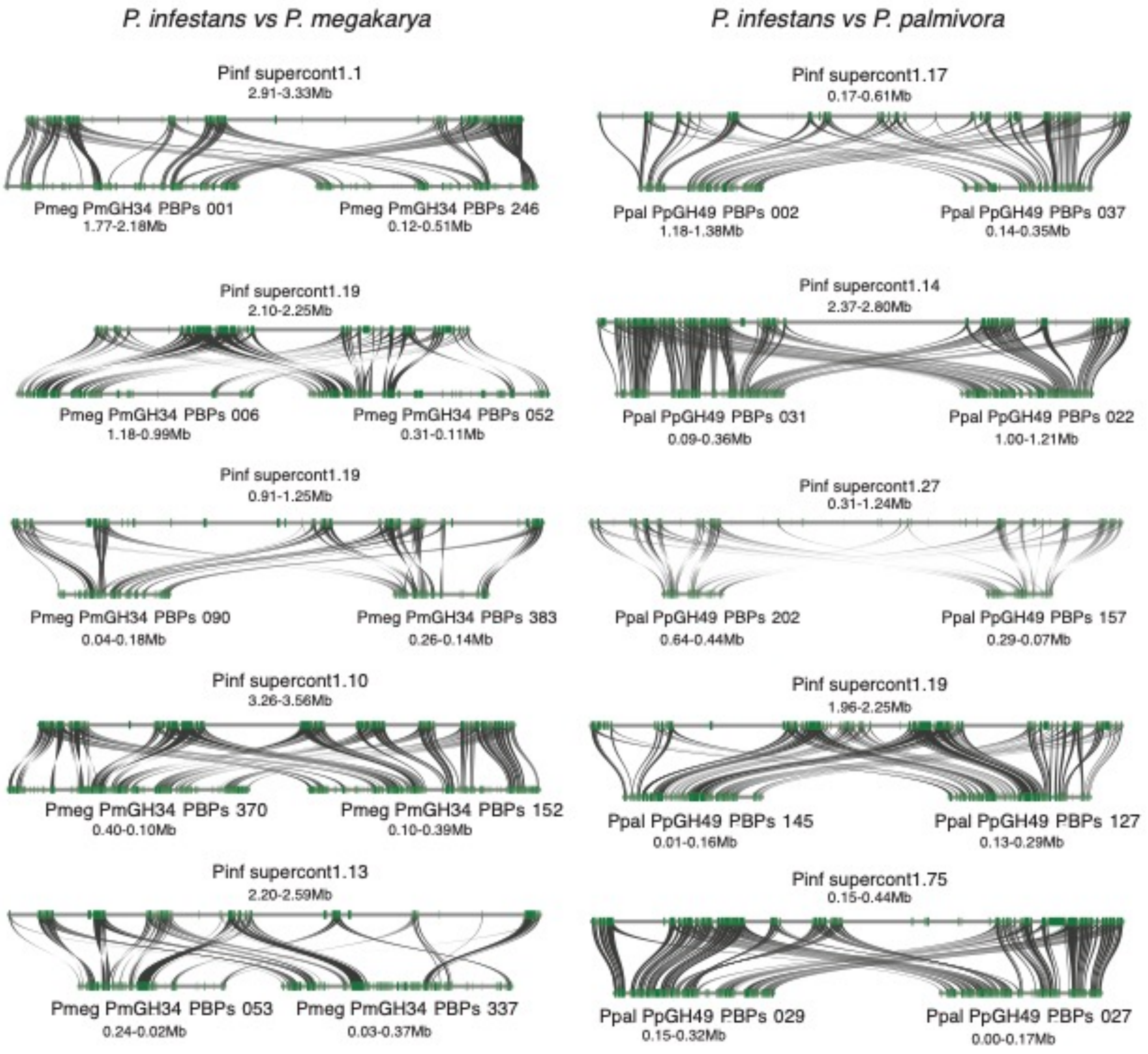


**Fig. S4.** Examples of duplicated regions in the genomes of *Pmeg* (Pm1/GH34) and *Ppal* (Pp2/GH49) syntenic to one region in the genome of *Pinf*.

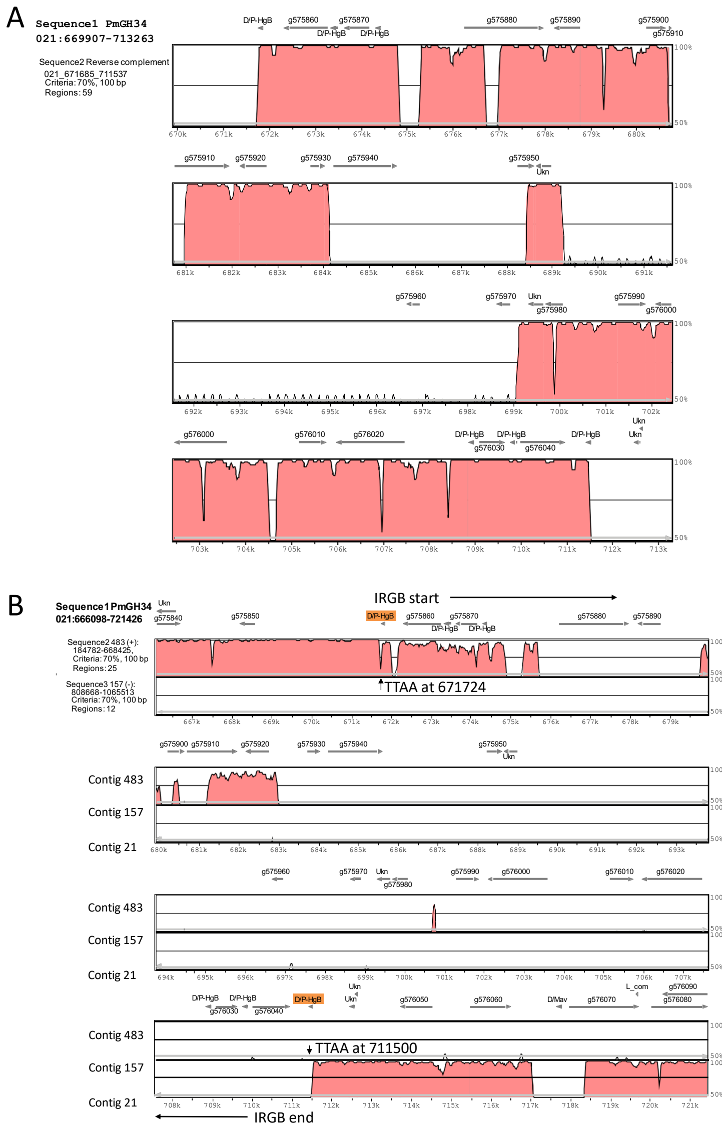


**Fig. S5.** *Pmeg* invert repeat gene block 21. (A) The invert repeat block on contig 21 aligned to its reverse complement showing inverted repeat segment on its end involving DNA/PiggyBac-like TEs (D/P-HGB) and multiple hypothetical protein encoding genes. (B) Invert repeat gene block 21 aligned to contigs 157 and 483 showing sequence homology with invert repeat gene block 21 flanking regions. TTAA sequence marks the ends of invert repeat gene block 21 and are shared by contigs 157 and 483 homologous regions.

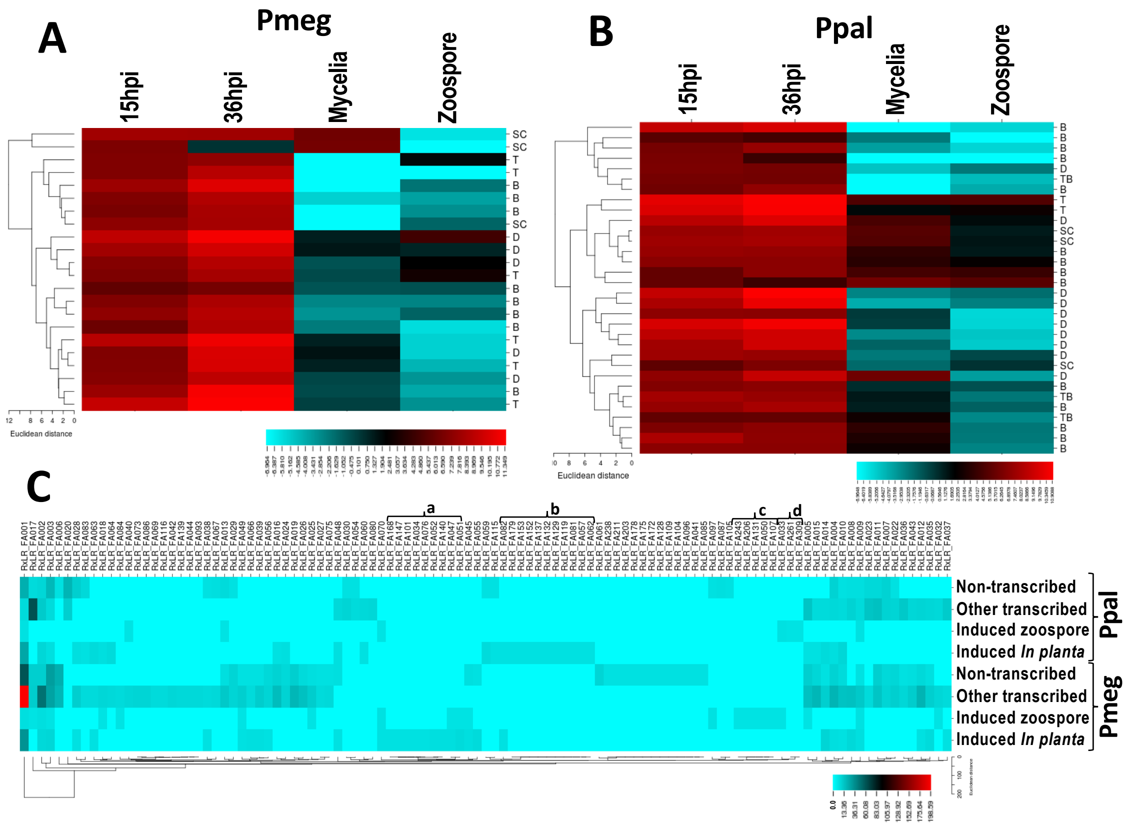


**Fig. S6.** Transcription profile of the *P. megakarya* (Pm1) and *P. palmivora* (Pp2) RxLRs detected *in planta* and distribution of the RxLR families based on their expression pattern. For the relative transcription profiles, *Pmeg* (A) and *Ppal* (B) RxLRs detected in at least three of the four biological replicates (with ≥1 paired read) of 15hpi *in planta* samples were normalized and $Log_{10}$ transformed. (C) Number of RxLRs in each of expression categories of the RxLR families. The heat map was generated using CIMminer (http://discover.nci.nih.gov/cimminer).

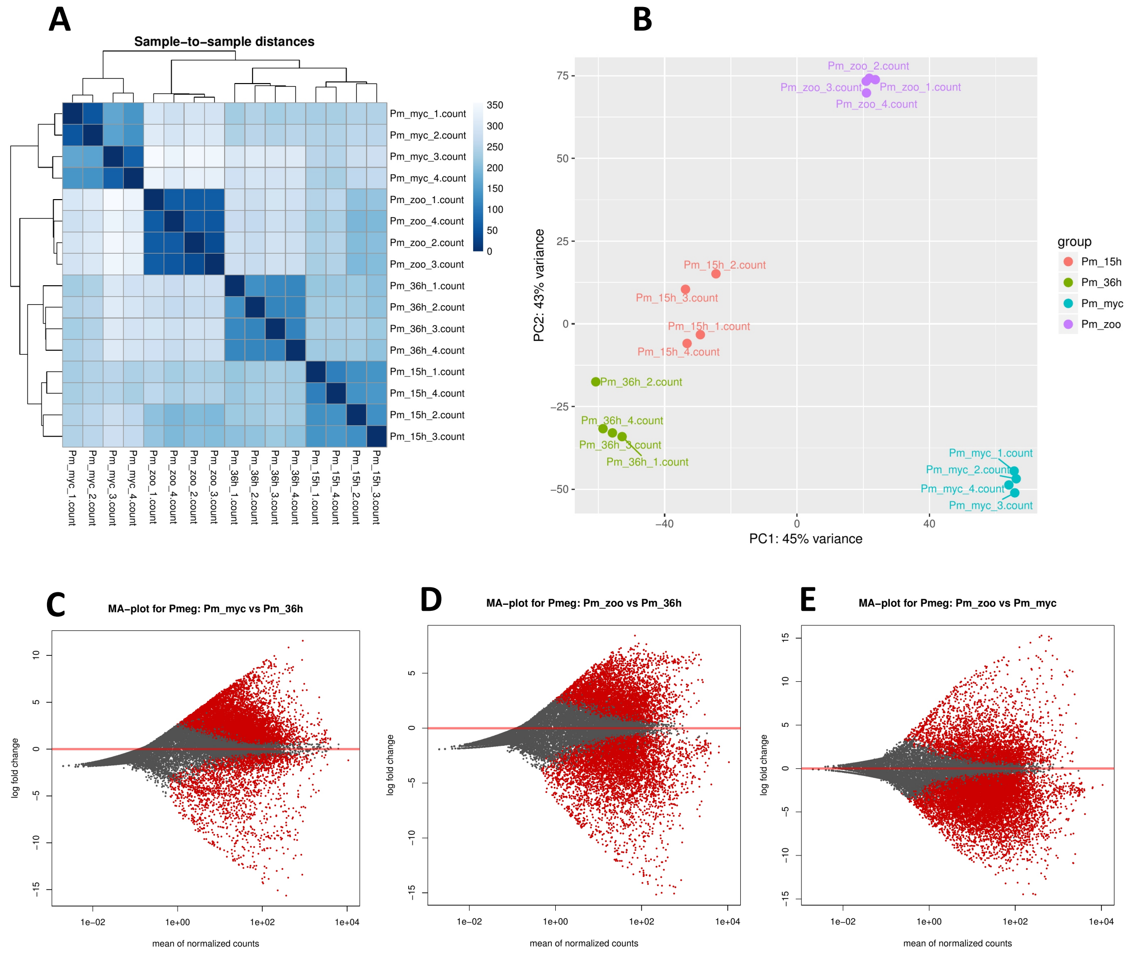


**Fig. S7.** RNASeq analysis revealed transcriptional difference in four different growth stages of *P. megakarya* (*Pmeg*) isolate Pm1. (A) Heatmap showing Euclidean distances among the 16 libraries, calculated from Variance stabilized transformed count data. Color key indicates level of similarity between libraries. (B) Principal Component Analysis (PCA) of the first two principal components (PC1, PC2) of the 16 libraries. (C-D) Scatter MA-plot showing differential expression for the *Pmeg* genes in mycelia, zoospore and in planta (36 h post inoculation). Red-colored dots represent DEGs with $log_2$-fold changes.

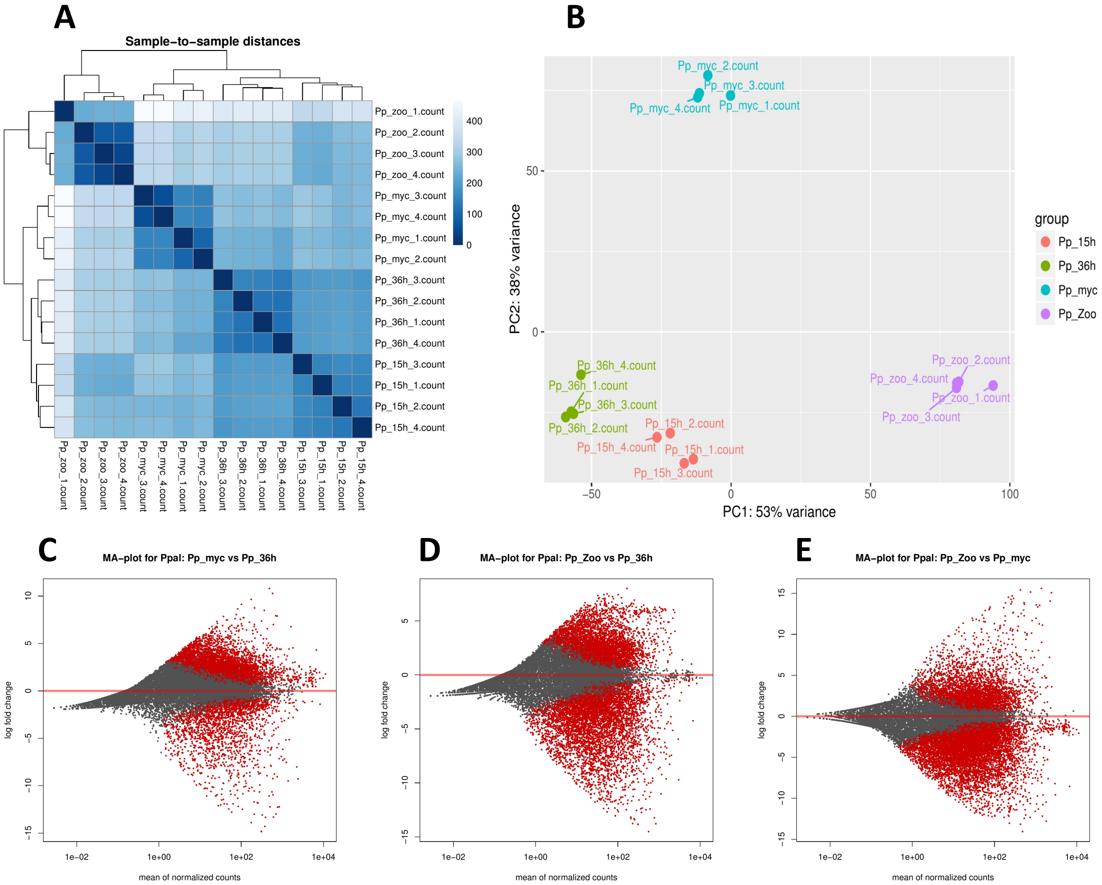


**Fig. S8.** RNASeq analysis revealed transcriptional difference in four different growth stages of *P. palmivora* (*Ppal*) isolate Pp2. (A) Heatmap showing Euclidean distances among the 16 libraries, calculated from Variance stabilized transformed count data. Color key indicates level of similarity between libraries. (B) Principal Component Analysis (PCA) of the first two principal components (PC1, PC2) of the 16 libraries. (C-D) Scatter MA-plot showing differential expression for the *Ppal* genes in mycelia, zoospore and in planta (36 h post inoculation). Red-colored dots represent DEGs with $log_2$-fold changes.

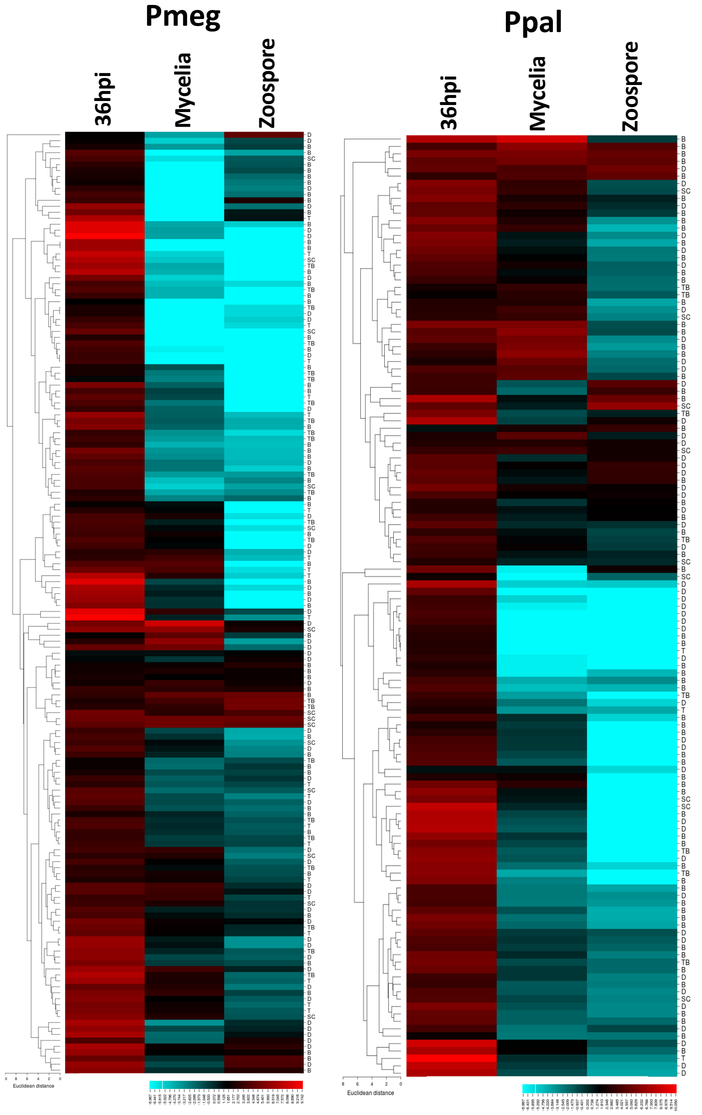


**Fig. S9.** Transcription profile of the *P. megakarya* (Pm1) and *P. palmivora* (Pp2) RxLRs detected *in planta*. For the relative transcription profiles, *Pmeg* (A) and *Ppal* (B) RxLRs detected in at least three of the four biological replicates (with ≥1 paired read) of 36hpi *in planta* samples were normalized and $Log_{10}$ transformed. The heat map was generated using CIMminer

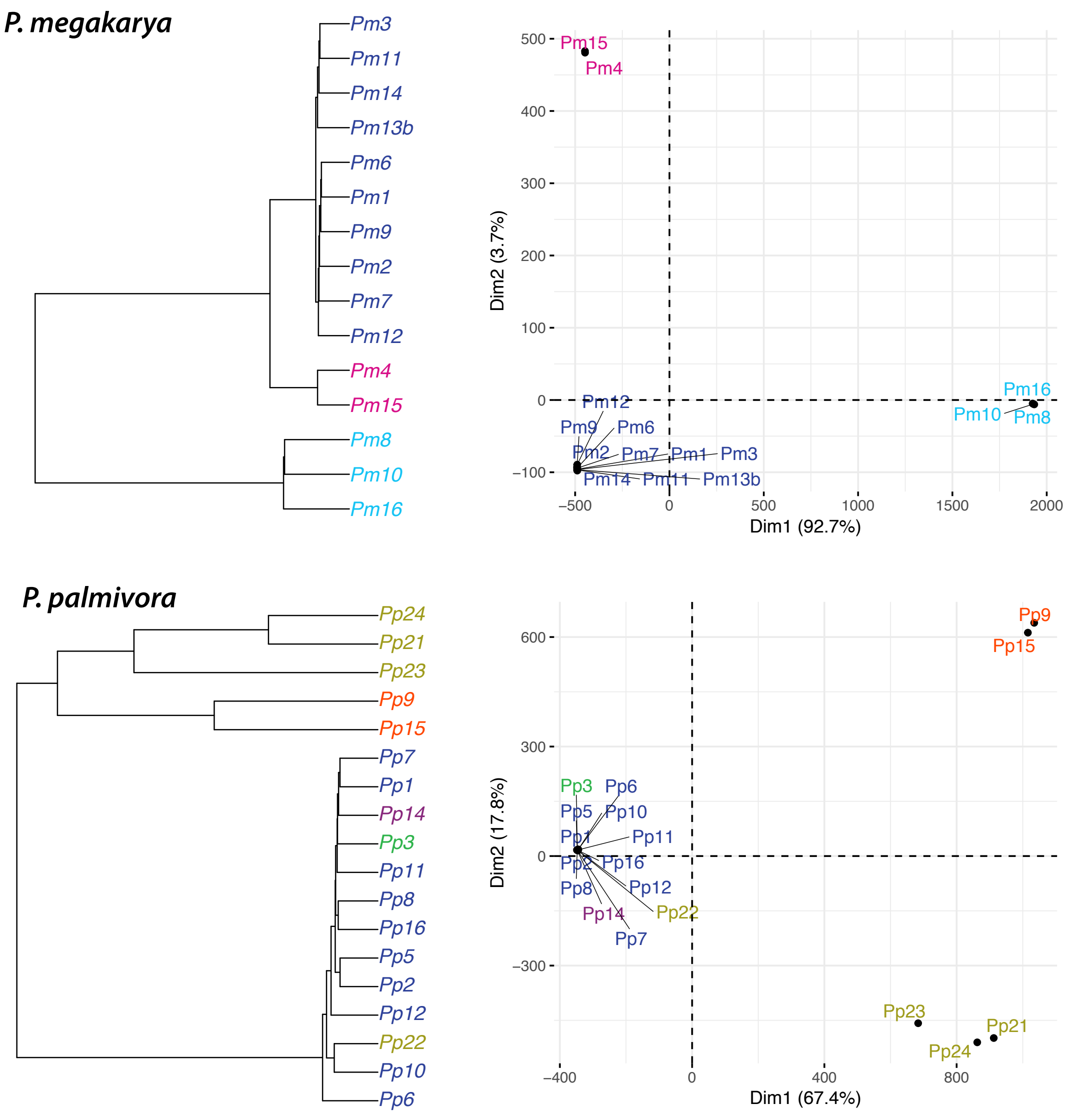


**Fig. S10.** Cladogram and PCA clustering re-sequenced isolates from the predicted SNPs. Isolates mostly clustered by their geographical origin (Green: Costa Rica, Purple: Côte d'Ivoire, Blue: Ghana, Magenta: Nigeria, Light blue: Cameroon, Orange: Indonesia and Papua New Guinea).

#### SUPPLEMENTARY TABLES

**Table S1.** Geographic origin of 34 Phytophthora isolates causing black pod of cacao.

| Isolate id | Isolate name | Species | Country of origin | Region/state | Year of isolation |
|---|---|---|---|---|---|
| Pm1 | GH34 | *P. megakarya* | Ghana | Eastern | 2013 |
| Pm10 | ZTHO131 | *P. megakarya* | Cameroon | Bakoa | 2011 |
| Pm11 | GH24 | *P. megakarya* | Ghana | Volta | 2013 |
| Pm12 | GH9 | *P. megakarya* | Ghana | Volta | 2013 |
| Pm13b | GH35 | *P. megakarya* | Ghana | Eastern | 2013 |
| Pm14 | GH41 | *P. megakarya* | Ghana | Eastern | 2013 |
| Pm15 | PH25 | *P. megakarya* | Nigeria | Ondo | 2016 |
| Pm16 | ZTHO144 | *P. megakarya* | Cameroon | Kedia | 2011 |
| Pm2 | GH12 | *P. megakarya* | Ghana | Volta | 2013 |
| Pm3 | GH10 | *P. megakarya* | Ghana | Volta | 2013 |
| Pm4 | PH95 | *P. megakarya* | Nigeria | Ondo | 2016 |
| Pm6 | GH7 | *P. megakarya* | Ghana | Volta | 2013 |
| Pm7 | GH19 | *P. megakarya* | Ghana | Volta | 2013 |
| Pm8 | ZTHO145 | *P. megakarya* | Cameroon | Kedia | 2011 |
| Pm9 | GH15 | *P. megakarya* | Ghana | Volta | 2013 |
| Pp1 | ER344 | *P. palmivora* | Ghana | Eastern | 2013 |
| Pp10 | WR411 | *P. palmivora* | Ghana | Western | 2012 |
| Pp11 | GH47 | *P. palmivora* | Ghana | Volta | 2013 |
| Pp12 | GH37 | *P. palmivora* | Ghana | Volta | 2013 |
| Pp14 | SBR112.9 | *P. palmivora* | Côte d'Ivoire | Bas-Sassandra | 2009 |
| Pp15 | PhAech | *P. palmivora* | Indonesia | Aceh | 2011 |
| Pp16 | AR61 | *P. palmivora* | Ghana | Ashanti | 1999 |
| Pp2 | GH49 | *P. palmivora* | Ghana | Eastern | 2013 |
| Pp3 | C18 | *P. palmivora* | Costa Rica | Caribic | 2014 |
| Pp5 | GH44 | *P. palmivora* | Ghana | Eastern | 2013 |
| Pp6 | WR417 | *P. palmivora* | Ghana | Western | 2012 |
| Pp7 | VR100 | *P. palmivora* | Ghana | Volta | 2012 |
| Pp8 | BAR133 | *P. palmivora* | Ghana | Brong Ahafo | 2011 |
| Pp9 | Asmn2 | *P. palmivora* | Indonesia | South Sulawesi | 2014 |
| Pp22 | COBP2 | *P. palmivora* | Papua New Guinea | East New Britain | 2005 |
| Pp23 | Mag40 | *P. palmivora* | Papua New Guinea | Madang | 2005 |
| Pp24 | NSP43 | *P. palmivora* | Papua New Guinea | Bougainville | 2005 |
| Pp21 | NSP61 | *P. palmivora* | Papua New Guinea | Bougainville | 2005 |

**Table S2.** Read statistics of long reads used for genome assemblies

| Metrics | Pm1/GH34 | Pm4 | Pm15 | Pp2/GH49 | Pp3 | Pp15 |
|---|---|---|---|---|---|---|
| Total number of bases (Kbp) | 29462.38 | 46174.80 | 23495.43 | 43036.97 | 37437.62 | 35842.18 |
| Number of reads | 2867546 | 3354633 | 2716479 | 4454033 | 2884835 | 2428092 |
| Average length of reads | 10274.4 | 13764.5 | 8649.22 | 9662.47 | 12977.4 | 14761.5 |
| Minimum length of reads | 50 | 50 | 50 | 50 | 50 | 50 |
| Maximum length of reads | 99988 | 113545 | 105072 | 79326 | 120334 | 128837 |
| Median length | 8706 | 10918 | 0.48 | 0.49 | 0.48 | 0.48 |
| N10 length | 26558 | 42903 | 28575 | 25213 | 38507 | 42429 |
| N20 length | 22659 | 34737 | 22982 | 21407 | 30916 | 34496 |
| N30 length | 20089 | 29306 | 19604 | 19035 | 26217 | 29493 |
| N40 length | 18096 | 25133 | 16949 | 17245 | 22809 | 25740 |
| N50 length | 16346 | 21701 | 14487 | 15591 | 20112 | 22668 |
| N60 length | 14458 | 18733 | 12059 | 13610 | 17756 | 19901 |
| N70 length | 12062 | 15768 | 9677 | 11240 | 15135 | 17070 |
| N80 length | 9277 | 12204 | 7275 | 8653 | 11700 | 13548 |
| N90 length | 6034 | 7876 | 4680 | 5665 | 7530 | 8714 |
| N10 index | 95668 | 89533 | 66779 | 146090 | 79898 | 70146 |
| N20 index | 216572 | 210063 | 159412 | 332679 | 189408 | 164617 |
| N30 index | 354981 | 355246 | 270492 | 546504 | 321414 | 277397 |
| N40 index | 509729 | 525728 | 399491 | 784347 | 474898 | 407743 |
| N50 index | 681010 | 723692 | 549338 | 1046599 | 649954 | 556290 |
| N60 index | 872151 | 952799 | 726843 | 1340974 | 847953 | 725017 |
| N70 index | 1094191 | 1220693 | 944052 | 1687945 | 1075330 | 918981 |
| N80 index | 1371166 | 1550992 | 1222991 | 2122097 | 1354559 | 1152625 |
| N90 index | 1759008 | 2015028 | 1620267 | 2727570 | 1747204 | 1476846 |
| Number of reads >= 100bp | 2857421 | 3327270 | 2688918 | 4433803 | 2865553 | 2416081 |
| Average length of reads >= 100bp | 10310.6 | 13877.1 | 8737.13 | 9706.21 | 13064.2 | 14834.5 |
| N50 reads >= 100bp | 16346 | 21701 | 14488 | 15591 | 20113 | 22669 |
| GC percentage | 0.5 | 0.48 | 6305 | 8089 | 10683 | 12573 |

**Table S3.** Genome assembly statistics by genome assembled in this study

| Metrics | Pm1/PmGH34 | Pm4 | Pm15 | Pp2/PpGH49 | Pp3 | Pp15 |
|---|---|---|---|---|---|---|
| Total number of bases | 214175571 | 250225369 | 201715625 | 115462317 | 144631407 | 145862580 |
| Number of contigs | 502 | 313 | 621 | 240 | 139 | 130 |
| Average length of contigs | 426645 | 799442 | 324824 | 481093 | 1040510 | 1122020 |
| Minimum length of contigs | 20151 | 2208 | 2834 | 22808 | 18022 | 4266 |
| Maximum length of contigs | 4094622 | 5827814 | 2651604 | 3040033 | 3667565 | 4740504 |
| N10 length | 2085467 | 3620582 | 1917074 | 2054601 | 3145829 | 3774787 |
| N20 length | 1674431 | 3029956 | 1327903 | 1468458 | 2693025 | 2959218 |
| N30 length | 1196352 | 2368594 | 1049413 | 1234300 | 2418398 | 2345091 |
| N40 length | 999003 | 1939916 | 814963 | 1074600 | 2174027 | 2012006 |
| N50 length | 780291 | 1658215 | 622146 | 877549 | 1822104 | 1877437 |
| N60 length | 620976 | 1323151 | 481591 | 689382 | 1409877 | 1576506 |
| N70 length | 451589 | 1019104 | 362673 | 499894 | 1147337 | 1405369 |
| N80 length | 342642 | 704814 | 255679 | 370881 | 827911 | 1047004 |
| N90 length | 214502 | 412506 | 158457 | 252349 | 650184 | 639164 |
| N10 index | 8 | 6 | 9 | 5 | 5 | 4 |
| N20 index | 19 | 13 | 22 | 12 | 10 | 8 |
| N30 index | 34 | 23 | 40 | 21 | 15 | 14 |
| N40 index | 54 | 35 | 62 | 31 | 22 | 20 |
| N50 index | 78 | 49 | 90 | 43 | 29 | 28 |
| N60 index | 109 | 66 | 127 | 58 | 38 | 36 |
| N70 index | 150 | 88 | 175 | 77 | 49 | 46 |
| N80 index | 204 | 118 | 242 | 104 | 64 | 57 |
| N90 index | 281 | 164 | 341 | 141 | 83 | 74 |
| Number of contigs >= 100bp | 502 | 313 | 621 | 240 | 139 | 130 |
| Average length of contigs >= 100bp | 426645 | 799442 | 324824 | 481093 | 1.04E+06 | 1.12E+06 |
| N50 contigs >= 100bp | 780291 | 1658215 | 622146 | 877549 | 1822104 | 1877437 |
| GC percentage | 49% | 50% | 49% | 49% | 49% | 49% |

**Table S4.** Summary table of the BUSCO genes detected per species.

| Species | Complete | Duplicated | Fragmented | Missing |
|---|---|---|---|---|
| *P. megakarya* | 0.86 | 0.32 | 0.026 | 0.1 |
| *P. palmivora* | 0.83 | 0.41 | 0.056 | 0.1 |
| *P. infestants* | 0.86 | 0.12 | 0.036 | 0.095 |
| *P. capsici* | 0.87 | 0.099 | 0.029 | 0.092 |
| *P. cinnamomi* | 0.85 | 0.11 | 0.049 | 0.099 |
| *P. ramorum* | 0.85 | 0.12 | 0.039 | 0.1 |
| *P. sojae* | 0.86 | 0.1 | 0.026 | 0.1 |

**Table S5.** Estimated haploid genome sizes (1C) estimated by flow cytometry.

| Species | Isolate | Replicates | Mean 1C (Mbp) | SD (Mbp) |
|---|---|---|---|---|
| *P. megakarya* | Pm1 | 3 | 231.47 | 0.97 |
| *P. megakarya* | Pm2 | 3 | 224.82 | 16.72 |
| *P. megakarya* | Pm4 | 3 | 241.92 | 6.36 |
| *P. megakarya* | Pm8 | 3 | 206.30 | 11.57 |
| *P. megakarya* | Pm15 | 3 | 220.28 | 8.63 |
| *P. palmivora* | Pp2 | 4 | 126.66 | 2.53 |
| *P. palmivora* | Pp3 | 3 | 132.98 | 3.92 |
| *P. palmivora* | Pp7 | 3 | 122.30 | 1.03 |
| *P. palmivora* | Pp14 | 3 | 121.05 | 3.30 |
| *P. palmivora* | Pp15 | 3 | 125.98 | 3.53 |

**Table S6.** General statistics of multiple assemblers

| Metric | Genotype | FALCON-Unzip | Canu | wtdbg2 |
| --- | --- | --- | --- | --- |
| Haploid assembly size (Mb) | *P. megakarya* Pm1 | 207 | 172* | 193 |
| Number of contigs | *P. megakarya* Pm1 | 1,163 | 2,205 | 2,729 |
| Maximum contig length (Mb) | *P. megakarya* Pm1 | 3.4 | 4.7 | 1.6 |
| N50 length (Kb) | *P. megakarya* Pm1 | 337 | 310 | 180 |
| Haploid assembly size (Mb) | *P. palmivora* Pp2 | 113 | 91* | 112 |
| Number of contigs | *P. palmivora* Pp2 | 484 | 886 | 2,358 |
| Maximum contig length (Mb) | *P. palmivora* Pp2 | 2.5 | 3.2 | 1 |
| N50 length (Kb) | *P. palmivora* Pp2 | 404 | 151 | 105 |

*Half of contig size

**Table S7.** Summary table from the repeat and transposable elements from *P. megakarya* and *P. palmivora* generated RepeatMasker.

| Class | Subclass | *P. megakarya* Pm1 | | | *P. palmivora* Pp2 | | |
|---|---|---|---|---|---|---|---|
| | | Count | bpMasked | %masked | Count | bpMasked | %masked |
| DNA | - | 1 | 201 | 0.00% | 26 | 2419 | 0.00% |
| | Academ-H | 48 | 6338 | 0.00% | - | - | - |
| | CMC-EnSpm | 539 | 117327 | 0.05% | 69 | 30222 | 0.03% |
| | Crypton | 1553 | 1297086 | 0.61% | 646 | 595064 | 0.52% |
| | Crypton-F | 648 | 237218 | 0.11% | 112 | 29792 | 0.03% |
| | Crypton-S | 1210 | 327654 | 0.15% | 813 | 234705 | 0.20% |
| | MULE-MuDR | 4567 | 1347121 | 0.63% | 2116 | 575225 | 0.50% |
| | Maverick | 511 | 123061 | 0.06% | 285 | 104153 | 0.09% |
| | Merlin | 59 | 18311 | 0.01% | 12 | 3691 | 0.00% |
| | MuLE-MuDR | 528 | 290288 | 0.14% | 403 | 277934 | 0.24% |
| | MuLE-NOF | 333 | 194421 | 0.09% | 60 | 81129 | 0.07% |
| | PIF-HarbS | 442 | 81338 | 0.04% | 278 | 45146 | 0.04% |
| | PIF-Harbinger | 1352 | 500090 | 0.23% | 182 | 82632 | 0.07% |
| | PiggyBac | 4141 | 2144246 | 1.00% | 2345 | 1092922 | 0.95% |
| | Sola | 241 | 76216 | 0.04% | 188 | 88659 | 0.08% |
| | Sola-1 | 322 | 91963 | 0.04% | 86 | 19683 | 0.02% |
| | Sola-3 | 127 | 71250 | 0.03% | 131 | 73537 | 0.06% |
| | TcMar-Ant1 | 201 | 137601 | 0.06% | 100 | 53334 | 0.05% |
| | TcMar-Fot1 | 40 | 22424 | 0.01% | 35 | 20446 | 0.02% |
| | TcMar-ISRm11 | 343 | 125457 | 0.06% | 312 | 109022 | 0.09% |
| | TcMar-Pogo | 354 | 63078 | 0.03% | 65 | 19299 | 0.02% |
| | TcMar-Stowaway | 219 | 64584 | 0.03% | 78 | 37791 | 0.03% |
| | TcMar-Tc1 | 160 | 74860 | 0.03% | 255 | 132874 | 0.12% |
| | TcMar-Tc2 | 658 | 266668 | 0.12% | 686 | 233533 | 0.20% |
| | hAT-Ac | 471 | 125794 | 0.06% | 409 | 87976 | 0.08% |
| | hAT-Charlie | 866 | 207129 | 0.10% | 3 | 2294 | 0.00% |
| | hAT-Tag1 | 235 | 111891 | 0.05% | 88 | 33160 | 0.03% |
| LINE | - | 43 | 9193 | 0.00% | 54 | 10927 | 0.01% |
| | CR1 | - | - | - | 1 | 49 | 0.00% |
| | I-Jockey | 108 | 36561 | 0.02% | 4 | 730 | 0.00% |
| | L1 | 1851 | 746208 | 0.35% | 587 | 181754 | 0.16% |
| | L1-Tx1 | 300 | 215898 | 0.10% | 58 | 58884 | 0.05% |
| | R1 | 95 | 14004 | 0.01% | 57 | 9207 | 0.01% |
| | R2 | 131 | 36765 | 0.02% | 20 | 6197 | 0.01% |
| | R2-NeSL | 242 | 363670 | 0.17% | 50 | 97381 | 0.08% |
| | RTE-BovB | 179 | 150630 | 0.07% | 73 | 33787 | 0.03% |
| LTR | - | 7 | 887 | 0.00% | 2 | 65 | 0.00% |
| | Copia | 8271 | 5568758 | 2.60% | 3135 | 2612083 | 2.26% |
| | ERV1 | 19 | 2179 | 0.00% | 13 | 1696 | 0.00% |
| | ERVL | 4 | 457 | 0.00% | 33 | 7590 | 0.01% |
| | Gypsy | 51960 | 52855979 | 24.68% | 22158 | 24255200 | 21.01% |
| | Gypsy-Cigr | 439 | 597592 | 0.28% | 303 | 295474 | 0.26% |
| | Ngaro | 1415 | 1071004 | 0.50% | 388 | 278393 | 0.24% |
| | Pao | 49 | 29956 | 0.01% | 16 | 7037 | 0.01% |
| RC | Helitron | 4445 | 3021199 | 1.41% | 1318 | 1120642 | 0.97% |
| SINE | - | 1775 | 437810 | 0.20% | 1423 | 297196 | 0.26% |
| | ID | 148 | 34091 | 0.02% | 152 | 49685 | 0.04% |
| | tRNA | 158 | 29159 | 0.01% | 168 | 17868 | 0.02% |
| Unknown | - | 18006 | 4597356 | 2.15% | 9378 | 2854466 | 2.47% |
| Low_complexity | - | 1005 | 52220 | 0.02% | 543 | 26319 | 0.02% |
| Satellite | - | 218 | 31344 | 0.01% | 73 | 9983 | 0.01% |
| Simple_repeat | - | 9807 | 622501 | 0.29% | 5324 | 285794 | 0.25% |
| rRNA | - | 168 | 143714 | 0.07% | 24 | 16389 | 0.01% |
| **Total** | | **121012** | **78792750** | **36.77%** | **55138** | **36601438** | **31.73%** |

**Table S8.** Transposable element count and mean size within 10 Kbp of the predicted genes in *P. megakarya.*

| Duplication pattern | Class of TE | Element count | Mean element size (bp) |
|---|---|---|---|
| BTD | LTR | 4.49 | 853.03 |
| DD | LTR | 7.41 | 991.75 |
| BD | LTR | 4.27 | 846.37 |
| TD | LTR | 6.17 | 949.12 |
| SC | LTR | 3.76 | 799.66 |
| BTD | DNA | 2.17 | 289.95 |
| DD | DNA | 2.60 | 338.43 |
| BD | DNA | 2.42 | 325.21 |
| TD | DNA | 2.41 | 334.67 |
| SC | DNA | 2.07 | 288.81 |
| BTD | LINE | 1.11 | 81.73 |
| DD | LINE | 1.17 | 131.71 |
| BD | LINE | 1.10 | 84.82 |
| TD | LINE | 1.16 | 145.78 |
| SC | LINE | 1.11 | 97.01 |
| BTD | SINE | 1.03 | 7.81 |
| DD | SINE | 1.08 | 19.23 |
| BD | SINE | 1.05 | 13.98 |
| TD | SINE | 1.03 | 6.12 |
| SC | SINE | 1.04 | 11.27 |
| BTD | RC | 4.41 | 203.01 |
| DD | RC | 3.06 | 186.02 |
| BD | RC | 3.32 | 178.12 |
| TD | RC | 3.36 | 216.53 |
| SC | RC | 3.19 | 163.82 |

**Table S9.** Potential infection-related genes in the *P. megakarya* and *P. palmivora* genomes

| **Gene classes** | ***P. megakarya*** | ***P. palmivora*** | ***P. sojae*** | ***P. parasitica*** | ***P. ramorum*** | ***P. infestans*** | ***P. capsici*** | ***P. cactorum*** |
|---|---|---|---|---|---|---|---|---|
| Proteases, all | 971(519) | 424(271) | 282 | 64 | 311 | nk | 40 | 87 |
| Serine proteases | 254(185) | 175(157) | 119 | 40 | 127 | 60 | 18 | 47 |
| Cysteine proteases | 220(45) | 59(22) | 67 | 24 | 74 | 33 | 22 | 40 |
| CAZymes, all | 730(632) | 611(563) | 388 | 514 | 324 | 435 | 628 | 901 |
| Carbohydrate esterases (CEs) | 58(50) | 36(33) | 37 | 46 | 21 | 49 | 86 | 111 |
| Glycoside hydrolases (GHs) | 403(346) | 332(307) | 196 | 280 | 167 | 244 | 261 | 374 |
| Glycosyl transferases (GTs) | 94(73) | 77(72) | 92 | 83 | 75 | 83 | 130 | 190 |
| Polysaccharide lyases (PLs) | 59(54) | 38(38) | 51 | 44 | 47 | 59 | 54 | 73 |
| Pectin lyases | 49(45) | 30(30) | 51 | 38 | 43 | 59 | 48 | 44 |
| Cutin hydrolase | 13(8) | 8(8) | 16 | 6 | 4 | 4 | 6 | 7 |
| Chitinases | 11(10) | 5(5) | 5 | 4 | 2 | nk | 2 | 3 |
| Protease inhibitors, all | 91(79) | 79(75) | 26 | 30 | 19 | 38 | 25 | 17 |
| Kazal | 63(55) | 63(59) | 15 | 28 | 12 | nk | 23 | 14 |
| Lipases, all | 112(88) | 99(98) | 171 | 59 | 154 | nk | 41 | 65 |
| NPP | 194(158) | 134(111) | 39 | 49 | 59 | 27 | 39 | 37 |
| *Effectors* | | | | | | | | |
| Elicitins | 69(65) | 68(62) | 57 | 54 | 50 | 40 | 48 | 39 |
| CRN | 197(156) | 177(127) | 41 | 13 | 19 | 196 | 25 | 16 |
| RxLR | 1381(1058) | 717(540) | 395 | 240 | 350 | 563 | 108 | 135 |
| *Detoxification genes* | | | | | | | | |
| ABC transporter | 279(114) | 124(90) | 141 | 48 | 135 | 137 | 40 | 60 |
| Cytochrome P450s | 89(71) | 20(14) | 31 | 40 | 31 | 28 | 36 | 46 |
| Alcohol dehydrogenase | 46(36) | 60(57) | 52 | 71 | nk | nk | 58 | 101 |
| Major Facilitator Superfamily (MFS) | 189(138) | 144(125) | 228 | 224 | nk | 111 | 217 | 239 |
| Short-chain dehydrogenase/reductase | 96(68) | 80(65) | 67 | 79 | nk | nk | 68 | 84 |

Number within parentheses indicates genes with >1 reads, either in mycelia, zoospores or in planta are considered transcribed.

**LEGENDS FOR DATASETS**

**Dataset S1.** Table with gene functional annotations of *P. megakarya* and *P. palmivora* and results from multiple analyses carried out in this study.

**Dataset S2.** Genomics coordinates of predicted repeats from *P. megakarya* and *P. palmivora.*

**Dataset S3.** Association of gene families and transposable elements.
